# A possible path to persistent re-entry waves at the outlet of the left pulmonary vein

**DOI:** 10.1101/2024.03.13.584755

**Authors:** Karoline Horgmo Jæger, Aslak Tveito

## Abstract

Atrial fibrillation (AF) is the most common form of cardiac arrhythmia, often evolving from paroxysmal episodes to persistent stages over an extended timeframe. While various factors contribute to this progression, the precise biophysical mechanisms driving it remain unclear. Here we explore how rapid firing of cardiomyocytes at the outlet of the pulmonary vein of the left atria can create a substrate for a persistent re-entry wave. This is grounded in a recently formulated mathematical model of the regulation of calcium ion channel density by intracellular calcium concentrations. According to the model, the density of membrane proteins carrying calcium ions is controlled by the intracellular calcium concentrations. In particular, if the concentration increases above a certain target level, the calcium current is weakened in order to restore the target level of calcium. During rapid pacing, the intracellular calcium concentration of the cardiomyocytes increases leading to a substantial reduction of the calcium current across the membrane of the myocytes, which again reduces the action potential duration. In a spatially resolved cell-based model of the outlet of the pulmonary vein of the left atria, we show that the reduced action potential duration can lead to re-entry.

Initiated by rapid pacing, often stemming from paroxysmal AF episodes lasting several days, the reduction in calcium current is a critical factor. Our findings illustrate how such episodes can foster a conducive environment for persistent AF through electrical remodeling, characterized by diminished calcium currents. This underscores the importance of promptly addressing early AF episodes to prevent their progression to chronic stages.

## 1 Introduction

Atrial fibrillation (AF) is the most common cardiac arrhythmia, characterized by rapid and irregular beating of the atria. This can lead to symptoms like palpitations, fatigue, and shortness of breath, increasing the risk of stroke and heart failure, [1, 2], along with a series of other possible health issues, [3]. The prevalence of AF is increasing and is expected to affect more than 8 million people in the USA by 2050, and more than 18 million in the EU by 2060, [4]. Indeed, significant societal and economic impacts of long-term AF are well-documented, [3, 2, 5]. While the biophysical mechanisms underlying AF have been studied for over a century, their precise nature remains a subject of ongoing debate, [6, 7, 8, 9, 10, 11]. Understanding of the mechanisms is crucial for effective treatment, including both ablation techniques and pharmacological interventions.

### Hypothesis

It appears to be a widespread consensus that AF typically progresses from paroxysmal episodes to persistent and eventually permanent forms [12, 10]. In this report, we explore a hypothesis aiming to explain the progression from paroxysmal to permanent atrial fibrillation (AF). Utilizing a recently developed mathematical model, we consider the potential impact of high-frequency firing over a limited period on the permanent establishment of re-entry, driven by remodeling of the calcium current in the cell membrane. The model suggests that increased intracellular calcium concentrations, triggered by high-frequency firing, lead to a reduction in the calcium current. This, in turn, shortens the action potential duration, thereby increasing the likelihood of re-entry. We present computational evidence to support the plausibility of this hypothesis, particularly focusing on the dynamics at the outlet of the left pulmonary vein.

The significance of the pulmonary veins’ outlets as primary sources of atrial fibrillation (AF) is well-established in the literature, [13, 14]. The prevalence of ectopic beats originating from this region is also well-documented, see, e.g., [15]. It is a recognized phenomenon that rapid firing of action potentials in the atria leads to an increase in intracellular calcium concentration, [16]. Furthermore, there is evidence suggesting that the calcium current is reduced during such rapid pacing, [17, 18, 19, 11], which in turn, is associated with a decrease in the duration of the action potential, [20].

When these observations are synthesized, the hypothesis presented above (see also Figure 1) is neither novel nor unexpected. However, the primary aim of this report is to demonstrate how these disparate elements coalesce within a computational model, thereby reinforcing the plausibility of the hypothesis.

**Figure 1:**
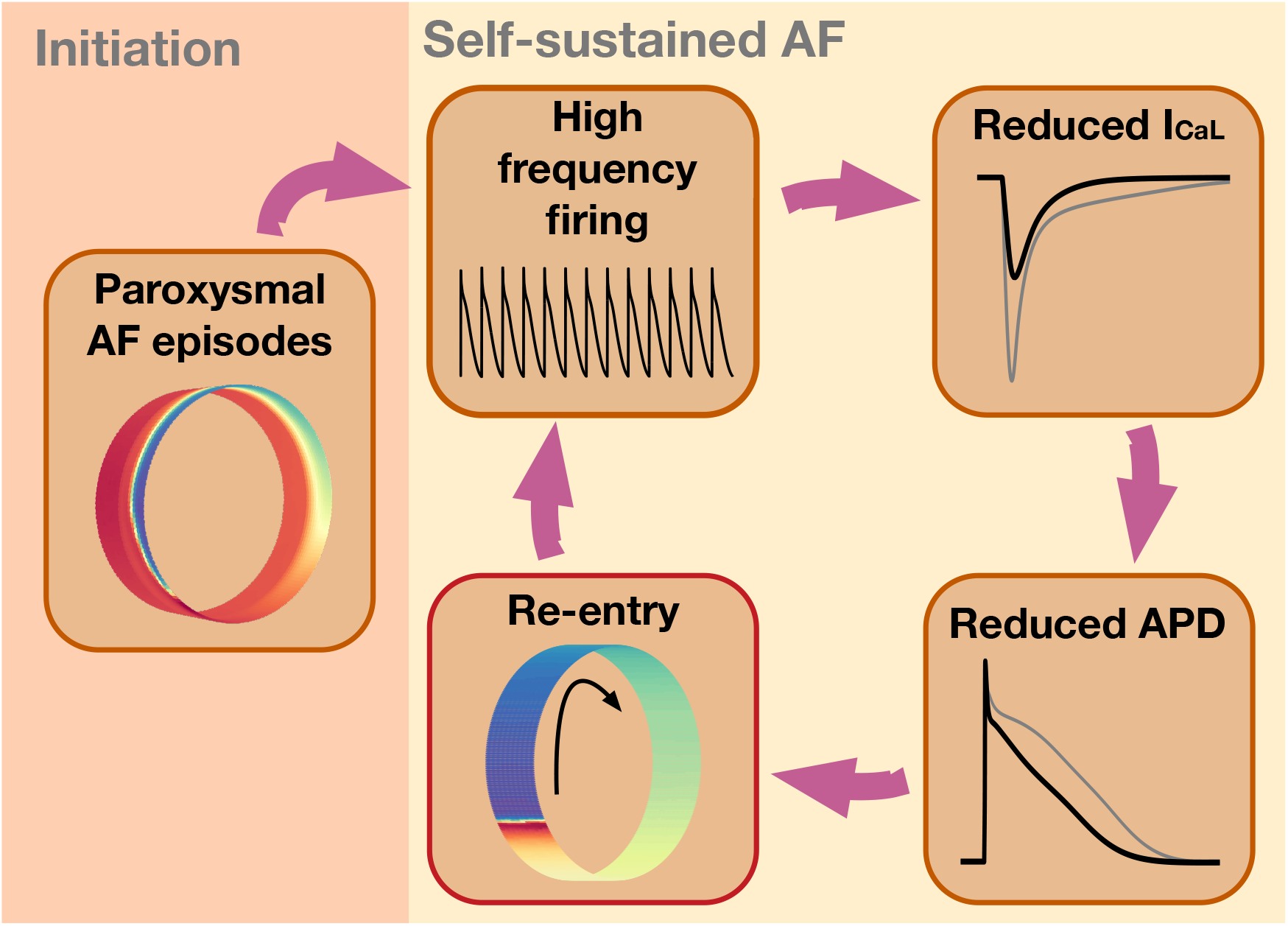
Hypothesis: The vicious cycle. Illustration of the proposed path to sustained re-entry around the PV sleeve. Paroxysmal AF episodes in the PV sleeve leads to a high frequency of action potential firing. According to the model (10), this leads to a reduction in the number of L-type calcium channels in the membrane of the cardiomyocytes in order to compensate for the increased intracellular calcium concentration resulting from rapid firing. The reduction in the number of L-type calcium channels reduces the L-type calcium current, *I*_CaL_. This reduction results in a reduction in the action potential duration (APD), which increases the likelihood of a re-entrant wave being able to travel around the PV sleeve. Such a re-entrant wave in itself results in a high firing frequency for the cardiomyocytes around the PV sleeve, which maintains the reduced *I*_CaL_ and reduced APD, promoting the continued existence of the re-entrant activity.

### Dynamics of the ion channel density of the *I*_CaL_-current

We will use a recently developed mathematical model to demonstrate a possible progressive path to sustained re-entry. In a series of papers, a theory on the regulation of the density of ion channels has been developed, see [21, 22, 23, 24, 25, 26, 27, 32]. For the regulation of the calcium current, the essence of this theory is that the density of ion channels is governed by the intracellular calcium concentration. If the intracellular calcium concentration, *c*, falls below a target value, *c*^∗^, the number of channels responsible for the *I*_CaL_-current is increased. The upregulation of the calcium current, *I*_CaL_, leads to a larger influx of Ca^2+^ into the cell and, as a result, more Ca^2+^ is released from internal storage systems because of the process referred to as graded release, see, e.g., [28, 29, 30]. The upregulation of *I*_CaL_ channels is continued until equilibrium is reached, i.e., until *c* = *c*^∗^. Conversely, if *c > c*^∗^, the number of ion channels is reduced until *c* is reduced to the target value. This model was developed for neurons [21, 22, 23, 24], and applied to study the dynamics of the Sino-atrial node [31]. Recently, the model was used to explain the time dependent efficacy of calcium channel blockers [32]. Note that for instance in [24] and [31] other currents and fluxes were also regulated by the level of intracellular calcium concentration, but we will concentrate on the *I*_CaL_ current for reasons that are explained in the Discussion.

### High frequency pacing increases intracellular calcium concentration

The hallmark of atrial fibrillation is an extremely high beat rate in the upper heart. This high frequency beating subsequently leads to increased *c* and this, in turn, according to the theory referred to above, down-regulates the *I*_CaL_-current. Reduced *I*_CaL_-current results in reduced action potential duration (APD) and this can be critical in order to maintain a re-entry wave. We will specifically show that sustained re-entry of an excitation wave around the pulmonary vein sleeve can be achieved in the following sequence 1) fast pacing by paroxysmal AF episodes, 2) down-regulation of the *I*_CaL_-current, 3) reduced APD, 4) high frequency pacing is self-sustained by the stable re-entry; adding up to a vicious cycle, see Figure 1.

## 2 Methods

In this section, we describe the models and methods used in the study. We describe the model used to represent the PV sleeve and the membrane models used to model the membrane dynamics of left atrial (LA) and pulmonary vein (PV) cardiomyocytes. In addition, we describe the model used for the regulation of the number of *I*_CaL_ channels. Finally, we present the numerical methods used in the simulations.

### 2.1 Cell-based mathematical models

In order to investigate the susceptibility of developing a re-entrant wave around the PV sleeve, we apply a cell-based mathematical model of the PV sleeve. We use the same set-up as that used to investigate the effect of mutations associated with AF in [33]. However, we will apply the more computationally efficient ‘simplified Kirchhoff’s network model’ (SKNM) [34] as opposed to the more detailed and considerably more computationally expensive extracellular-membrane-intracellular (EMI) model (see, e.g., [35, 36, 37, 38, 39, 40]) applied in [33]. The SKNM is a simplification of Kirchhoff’s network model (KNM), and KNM has been shown to provide good approximations of the EMI model in simulations of cardiac tissue [41].

The SKNM is derived from KNM based on the same simplifying assumption as that used to derive the monodomain model from the bidomain model of cardiac electrophysiology, and it has been shown that SKNM in several cases is a good approximation of KNM, see [34]. In the Supplementary Information, we compare the solution of KNM and SKNM for some simulations of the PV sleeve, and we confirm that the solutions of the two models are similar also in this case (see Figure S2).

As mentioned above, the change of model from EMI to SKNM results in a considerable reduction in the required computational efforts. In [33], a 500 ms EMI model simulation of the PV sleeve required a simulation time of about 8 days. Using SKNM, we are in this study able to perform the same simulation in about 5 minutes using the same computer. For comparison, the corresponding KNM simulations required a simulation time of about 6 hours. The reduced CPU efforts are important since they allow for large numbers of simulations, which will be necessary in order to investigate under what conditions sustainable re-entry will appear.

#### 2.1.1 SKNM equations

The SKNM equations are given by

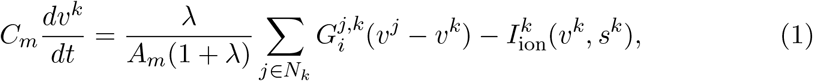

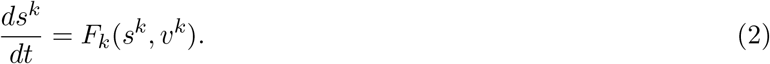

Here, *C*_*m*_ is the specific membrane capacitance (in *µ*F/cm^2^), *A*_*m*_ is the membrane area of one cell (in cm^2^), *v*_*k*_ is the membrane potential of cell *k* (in mV), *N*_*k*_ is the collection of all the neighbors of cell *k*, 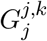 is the intracellular conductance between cells *k* and *j* (in mS), and *λ* is a scaling factor involved in the derivation of SKNM from KNM (see [34]). This parameter will be defined below. Moreover, 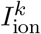 is the current density though ion channels, pumps and exchangers in the membrane of cell *k* (in *µ*A/cm^2^), *s*^*k*^ is a set of additional state variables involved in the computation of 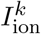, including gating variables for the ion channels and ionic concentrations, and *F*_*k*_ govern the dynamics of these variables. The models used for 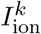, *s*^*k*^ and *F*_*k*_ are described in Section 2.2, and the details of the membrane model is provided in the Supplementary Information.

#### 2.1.2 SKNM parameters

The default parameters used in the SKNM simulations of this study are given in Table 1. The SKNM parameters 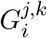 and *λ* are defined as described in [34]. In short,

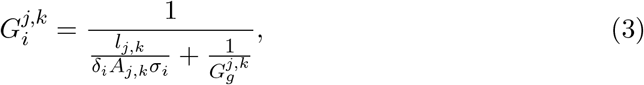

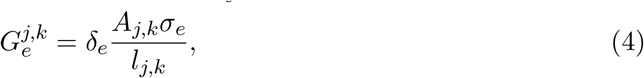

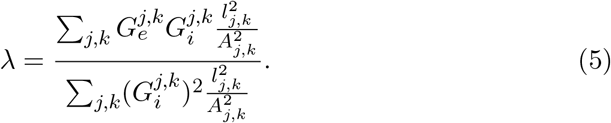

Here, *l*_*j,k*_ is the distance between the centers of cells *k* and *j* (in cm), *δ*_*e*_ (unitless) is the average extracellular volume fraction, *δ*_*i*_ = 1 − *δ*_*e*_ (unitless) is the average intracellular volume fraction, *A*_*j,k*_ (in cm^2^) is the average cross-sectional areas of compartments *j* and *k* (containing both intracellular and extracellular space), *σ*_*i*_ and *σ*_*e*_ are the intracellular and extracellular conductivities, respectively, (in mS/cm), 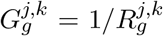 is the conductance of the gap junctions connecting cells *j* and *k* (in mS), and 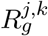 is the corresponding gap junction resistance (in kΩ).

**Table 1:** Default parameter values used in the SKNM simulations, based on [33]. Here, *l*_*l*_ is the cell length in the longitudinal direction and *l*_*t*_ is the maximal cell width in the transverse direction. Similarly, 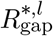 and 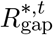 are the default gap junction resistances in the longitudinal and transverse directions, respectively.

| Parameter | Value |
| --- | --- |
| $C_m$ | $1 \mu\text{F}/\text{cm}^2$ |
| $A_m$ | $5 \cdot 10^{-5} \text{ cm}^2$ |
| $\sigma_i$ | $4 \text{ mS}/\text{cm}$ |
| $\sigma_e$ | $20 \text{ mS}/\text{cm}$ |
| $\delta_e$ | $0.2$ |
| $l_l$ | $120 \mu\text{m}$ |
| $l_t$ | $14 \mu\text{m}$ |
| $\lambda$ | $3.25$ |
| $R_{\text{gap}}^{*,l}$ | $380 \text{ k}\Omega$ |
| $R_{\text{gap}}^{*,t}$ | $760 \text{ k}\Omega$ |

To represent the slow and complex conduction conduction observed in the PV sleeve [42], we vary the value of the gap junction resistance randomly for each cell connection like in [33]. More specifically, for each cell connection, we draw a random number, *β*^*j,k*^, between 0 and 1 and let the gap junction resistance be given by

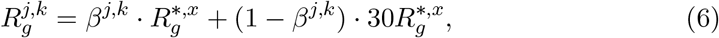

where 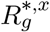 is either 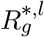 or 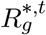 specified in Table 1, depending on whether the connection between cells *j* and *k* is in the longitudinal or transverse cell directions.

#### 2.1.3 The mesh defined by cardiomyocytes

In SKNM, the location of the cells define the computational mesh. In our simulations, we consider a collection of cells located in a cylinder at the outlet of a pulmonary vein. We use the same PV sleeve geometry as in [33]. That is, each cell is assumed to have a maximal diameter of 14 *µ*m and a length of 120 *µ*m, and the cells are connected in the longitudinal direction around the cylinder. In addition, 10 rows of connected cells are located along the longitudinal axis of the cylinder. In the default case, the PV sleeve is assumed to have a diameter of 1.5 cm, similar to diameters measured for human pulmonary veins [43]. This diameter corresponds to 393 cell lengths. Thus, in the default case, the domain consists of 393*×*10 cells, see leftmost panel of Figure 2.

**Figure 2:**
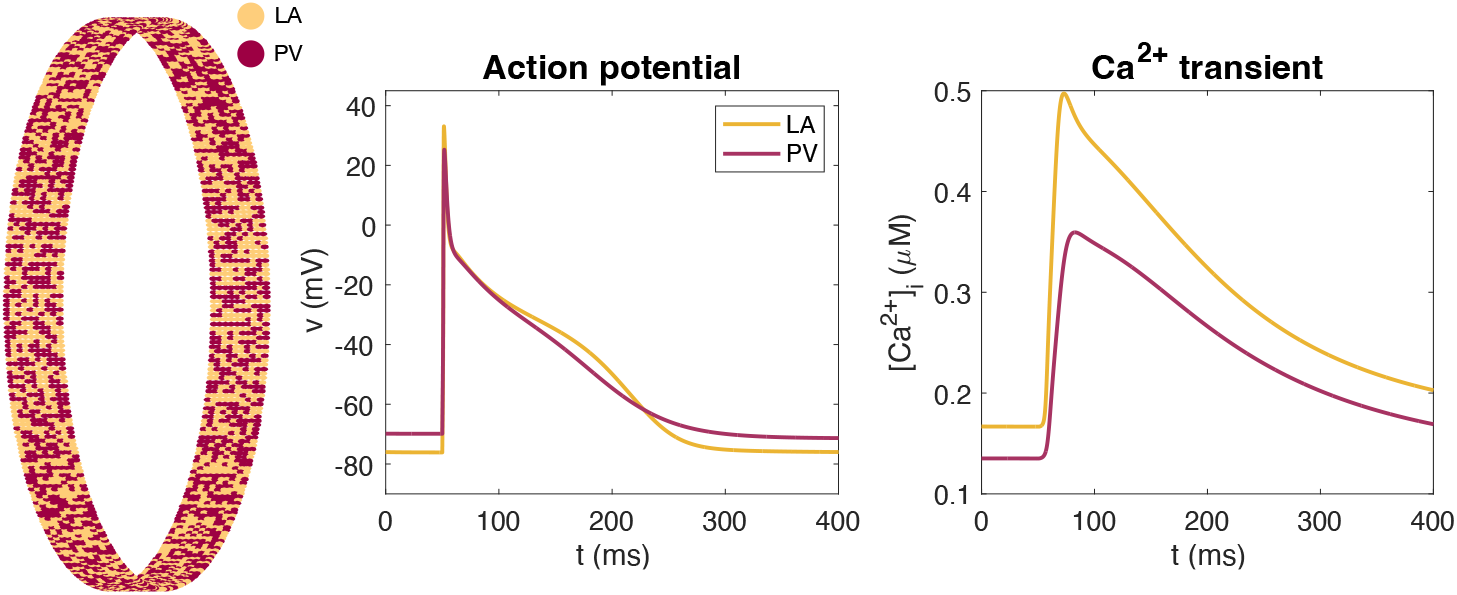
Mix of PV and LA cardiomyocytes. Illustration of the mix of LA and PV cardiomyocytes in the SKNM representation of the PV sleeve. The left panel shows an overview of the location of LA and PV myocytes in the PV sleeve. The density of certain ion channels are different for the PV and LA myocytes, resulting in the displayed differences in the action potential and calcium transients in single-cell simulations of the two cell types, shown in the center and right panels. The full model formulation is found in the Supplementary Information.

### 2.2 The PV-LA cell-based model

In the SKNM collection of cells, we assume that the cells are either ordinary left atrial (LA) cardiomyocytes, or pulmonary vein (PV) cardiomyocytes. These two cell types have been shown to have some differences in the density of some types of ion channels [44]. The differences are incorporated into the model used to define the membrane dynamics, 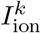 and *F*_*k*_. We use the membrane models from [33] with a few parameter adjustments to represent the two cell types. These adjustments will be described in Section 2.4 below. In addition, the details of the membrane models are provided in the Supplementary Information. In Figure 2 we show the distribution of cell types in the PV sleeve cylinder and the action potential and calcium transients of the two versions of the membrane model. In the default case, we consider 50% PV cells and 50% LA cells.

### 2.3 Modeling the expression of calcium channels

A model of the expression of ion channels carrying the *I*_CaL_ current can be written on the form

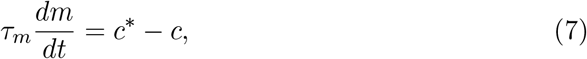

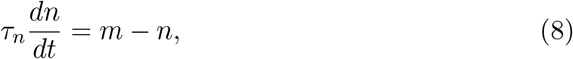

see [24, 31, 32]. In this model, *τ*_*m*_ (in mMms) and *τ*_*n*_ (in ms) are time constants, *c* (in mM) is the cytosolic calcium concentration, and *c*^∗^ (in mM) is the target level of the cytosolic calcium concentration. Furthermore, *m* = *m*(*t*) and *n* = *n*(*t*) denote the relative changes of the number of the messenger RNAs, *M* = *M* (*t*) and the number of expressed calcium ion channel proteins in the cell membrane, *N* = *N* (*t*), respectively. Specifically, we let *M*_0_ and *N*_0_ denote the default values of *M* and *N*, and therefore *M* (*t*) = *m*(*t*)*M*_0_ and *N* (*t*) = *n*(*t*)*N*_0_. The number of ion channels carrying the *I*_CaL_-current affects the strength of the current as follows,

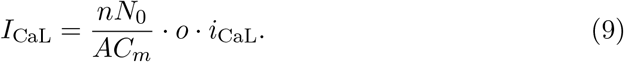

Here, *C*_*m*_ is the specific membrane capacitance (in *µ*F/cm^2^), *A* is the area of the cell membrane (in cm^2^), *o* is the (unitless) open probability, and *i*_CaL_ is the average current through a single open calcium channel (in *µ*A).

One disadvantage of this model is that it does not impose a natural upper or lower bound on the number of ion channels. In [32] it was shown that the model (7)– (8) can be accurately approximated by the following scalar model which includes bounds on the number of ion channels,

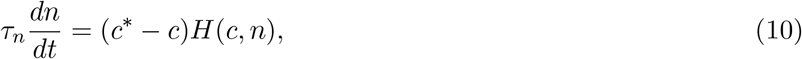

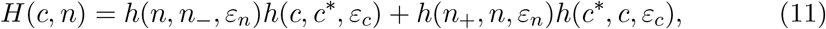

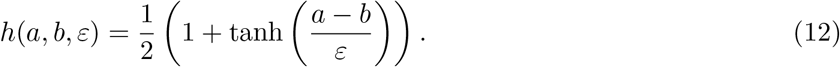

Here, *n* = *n*_−_ and *n* = *n*_+_ defines lower and upper bounds for the relative changes in the number of ion channels *N* = *N* (*t*). A plot of the function *H* are given in the Supplementary Information. In our computations, we use the model (10)–(12) to model the regulation of the number of *I*_CaL_ channels.

### 2.4 Parameterization of the model for channel expression

In the computations presented here, we have set the lower bound of channels to *n*_−_ = 0.28, motivated by the fact that *I*_CaL_ has been observed to be reduced to about 28% in patients with persistent AF [45]. In order to achieve a value of *n* close to this experimental value in the LA model following 600 beats per minute (bpm) pacing, we also needed to make some adjustments of the membrane model parameterizations. Specifically, *Ī*_NaCa_ is reduced by 40%, *g*_Kr_ is reduced by 20% and the calcium buffer concentrations 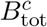 and 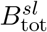 are both increased by 40% in the updated versions of the models.

For the remaining channel expression model parameterization, we have re-used the parameter *n*_+_ = 3 from [32], but since *n* is decreasing and not increasing in our simulations, this value does not influence the results of this study. Furthermore, the value of *τ*_*n*_ is set to 30,000 mMms. This value was chosen because it resulted in a reduction of *I*_CaL_ of about 50% after approximately 12 hours of rapid pacing (600 bpm) in the left atrial (LA) membrane model, consistent with experimental measurements of *I*_CaL_ in rabbit right atria at 600 bpm pacing from [17]. The value of *c*^∗^ for LA and PV is defined as the average value of the cytosolic calcium concentration over one second in the default versions of the LA and PV membrane models paced at 1 beat per second (1 bps).

### 2.5 Numerical implementations

We perform simulations using SKNM (1)–(2) coupled to the membrane models described in Section 2.2 as well as simulations of the single-cell membrane models in the form

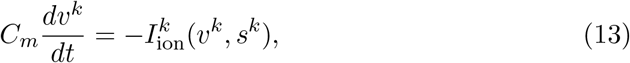

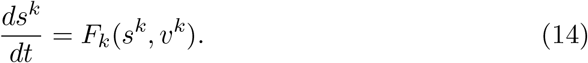

The single-cell membrane model simulations are performed in MATLAB using the ode15s solver. The SKNM simulations are done in C++ using a standard operator splitting of the linear and non-linear systems (see, e.g., [46, 47]) and a time step of Δ*t* = 0.001 ms. The linear part of the system is solved implicitly using the BiCGSTAB iterative solver from the MFEM library [48, 49], and the non-linear part is solved using the first-order Rush-Larsen method [50, 51] with code generated by the Gotran code generator [52]. We apply OpenMP parallelization [53] for the solution of the non-linear equations. The computations are run on a Dell Precision 3640 Tower with an Intel Core processor (i9-10900K, 3.7 GHz/5.4 GHz) with 10 cores, each with 2 threads.

## 3 Results

In this section, we present results of numerical simulations. First, we confirm the assumption that a reduction in *I*_CaL_ promotes the existence of a re-entrant wave around the computational PV sleeve. Next, we confirm that rapid pacing resulting from paroxysmal AF leads to a reduction in *I*_CaL_ in the LA and PV membrane models with the model (10) for the expression of *I*_CaL_ channels. This reduction in *I*_CaL_ is next shown to, in some cases, be sufficient for a re-entrant wave around the PV sleeve to be generated. Such a re-entrant wave would then in itself be responsible for rapid pacing and a reduced *I*_CaL_, making the mechanism self-sustained (see Figure 1). Finally, we show how perturbations of physiological parameters influence the forming of re-entry waves.

### 3.1 Reduced *I*_CaL_ promotes re-entry around the PV sleeve

In order to investigate re-entry susceptibility, we use the same set-up as in [33], illustrated in Figure 4B. The upper panel of Figure 3 shows the solution of such a simulation in the default case. We observe that following a stimulation (e.g., an ectopic beat), an excitation wave is generated traveling around the PV sleeve, but when the wave has traveled one round around the cylinder, the initially stimulated cells are not sufficiently repolarized to be excited again and the wave terminates after traveling one round.

**Figure 3:**
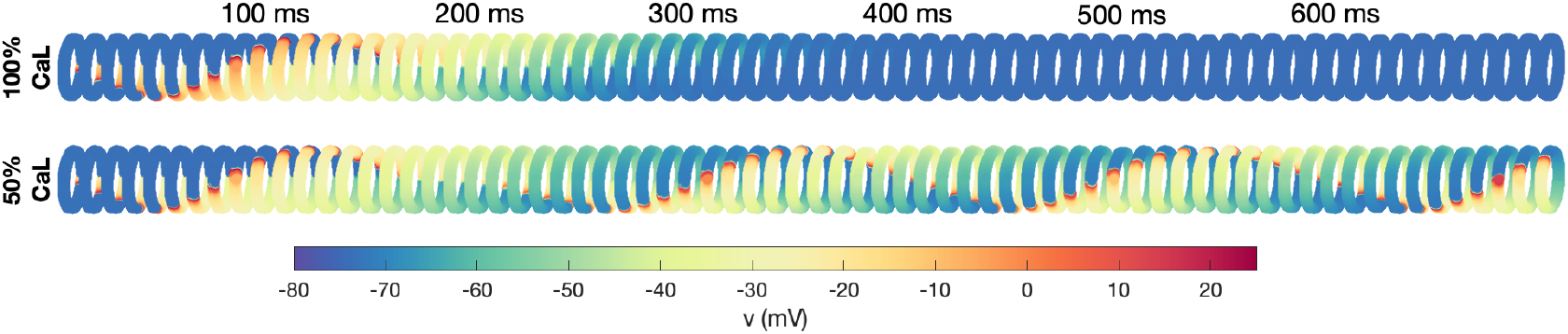
Re-entry around the PV sleeve for reduced *I*_CaL_. The figure shows the membrane potential for the cells in the PV sleeve cylinder at some different points in time after a stimulation is applied (see Figure 4B), representing an ectopic beat. The upper panel shows the solution in the default case, and the lower panel shows the solution when the *I*_CaL_-current has been reduced by 50%, which induces sustained re-entry.

**Figure 4:**
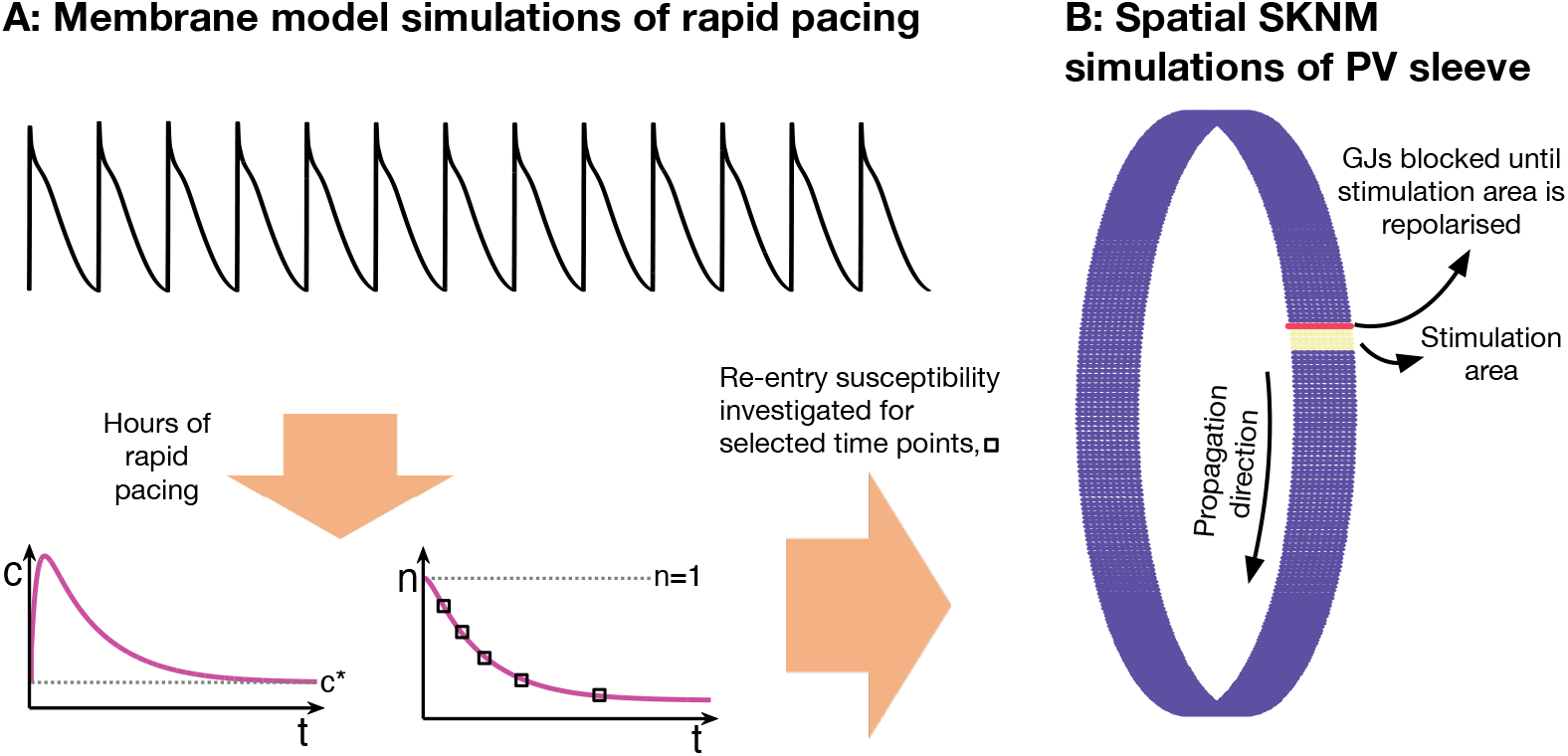
Computational procedure. Illustration of the computational procedure used to investigate re-entry susceptibility in the PV sleeve. **A:** First, membrane model simulations of the LA and PV cardiomyocytes with rapid pacing are conducted. After minutes and hours of rapid pacing, the number of *I*_CaL_ channels, *n*, is considerably reduced to compensate for the increased intracellular calcium concentration resulting from the rapid pacing. **B:** At some selected points in time, *n* is collected from the membrane model simulations and an SKNM simulation is conducted to investigate if a re-entrant wave traveling around the PV sleeve can be generated. In order to generate a wave traveling in one direction, the cells in a small area of the PV sleeve are stimulated and the gap junction (GJ) connections on the boundary of this area in one of the directions are blocked until the stimulated cells are repolarized.

In the lower panel of Figure 3, however, we consider the case of *I*_CaL_ reduced by 50%. In this case, the cells repolarize faster (shorter action potential duration) and the initially stimulated cells are ready to be excited again when the excitation wave has traveled one round. Consequently, the wave continues to continuously travel around and around the PV sleeve. The wave travels with a conduction velocity of about 29.6 cm/s. This means that the wave uses about 160 ms to travel around the cylinder of diameter 1.5 cm, corresponding to a beat rate of about 6.3 beats per second (bps) or 375 beats per minute (bpm) for the PV sleeve cardiomyocytes.

### 3.2 A computational path to sustained re-entry

Figure 3 illustrates that a reduction in the *I*_CaL_-current might promote the generation of a re-entrant wave around the PV sleeve resulting in a high beat rate. The purpose of this paper is to illustrate that such a reduction in the *I*_CaL_-current might result from rapid firing because rapid firing increases the intracellular calcium concentration, which according to the model (10) would result in a reduction in the number of *I*_CaL_-channels, *n*. As illustrated in Figure 1, once a re-entrant wave is generated, this re-entrant wave would result in rapid firing of the cells which could maintain the reduced number of *I*_CaL_ channels and increased re-entry susceptibility in a self-sustained manner. However, in order for the re-entrant wave to be generated in the first place, some other mechanism (e.g., paroxysmal AF episodes, see Figure 1) is needed to produce the initial rapid beating resulting in the initial reduction of the number of *I*_CaL_-channels.

We investigate this mechanism using the set-up described in Figure 4. First, we perform membrane model simulations of rapid pacing and observe how the number of *I*_CaL_-channels, *n*, is reduced over time. For a number of selected time points, we then investigate whether the *I*_CaL_ reduction at that point in time is enough for a re-entrant wave to be generated using the SKNM representation of the PV sleeve.

### 3.3 Development of the *I*_CaL_ current under influence of rapid firing

In Figure 5, we show the solution of an LA membrane model simulation with a pacing of 10 bps (or 600 bpm). We observe that the intracellular calcium concentration increases as a result of the rapid pacing and that the number of *I*_CaL_ channels consequently is reduced. After about 12 h, the number of *I*_CaL_ channels is reduced to about 50% (*n* = 0.5) and after about 150 h, the number of channels is reduced to about 32% (*n* = 0.32). The right panel illustrates *I*_CaL_ and the action potential as some selected points in time during the simulation. We observe that the action potential duration is reduced as the rapid pacing continues.

**Figure 5:**
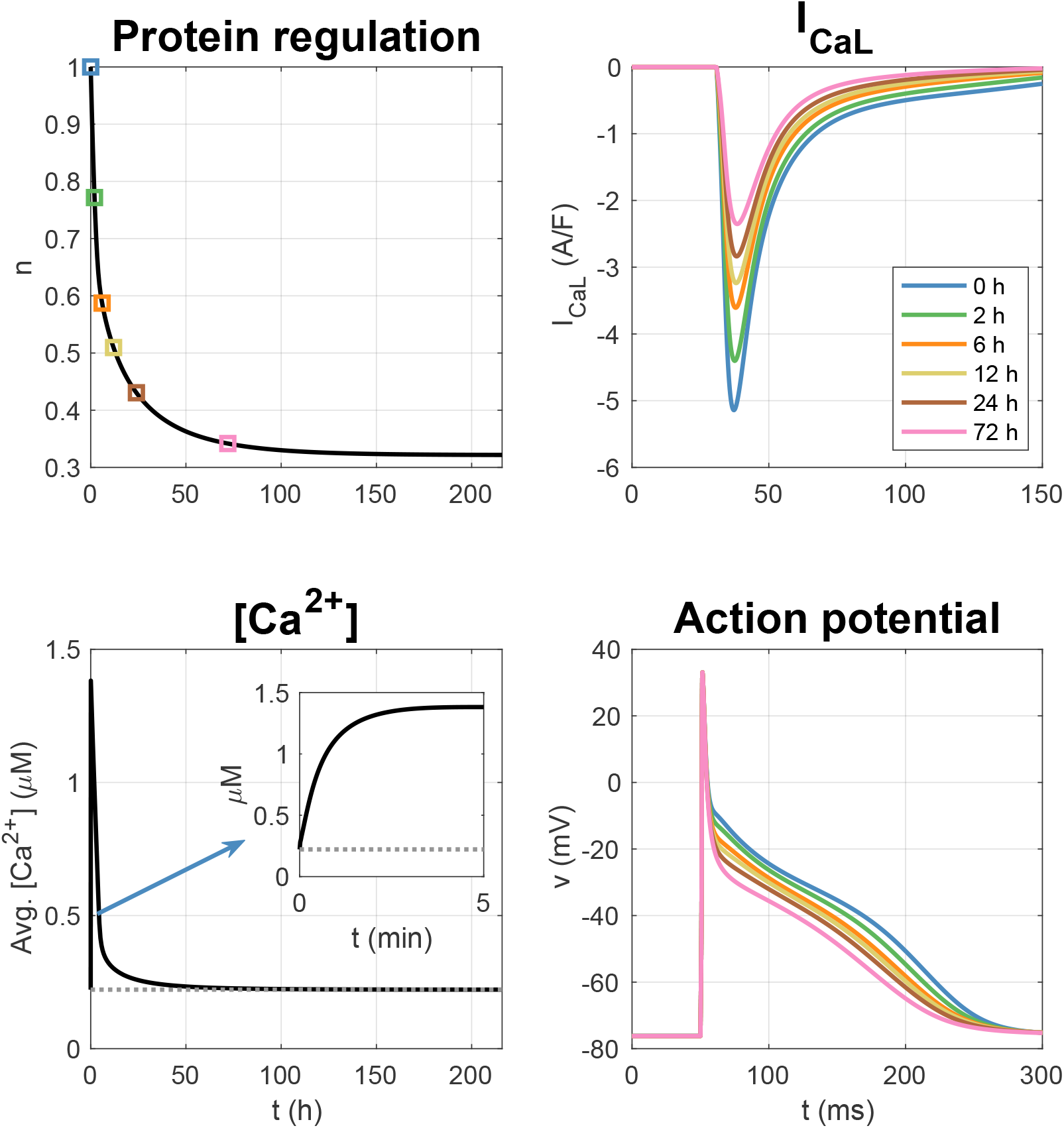
LA membrane model simulation with pacing of 10 bps. The upper left panel shows how the relative number of *I*_CaL_ channels, *n*, is reduced in response to the increased intracellular calcium concentration resulting from the rapid pacing. The lower left panel shows the average intracellular calcium concentration (averaged over a second) as a function of time. The gray dotted line is the target calcium concentration, *c*^∗^. The upper right panel shows *I*_CaL_ at given points in time during the simulation, and the lower right panel shows the corresponding action potentials.

In Figure 6, we show the solution of similar simulations using different pacing frequencies for both the LA and PV membrane models. We observe that the reduction in the number of *I*_CaL_ channels and the reduction in the action potential duration is faster and more pronounced as the pacing frequency increases.

**Figure 6:**
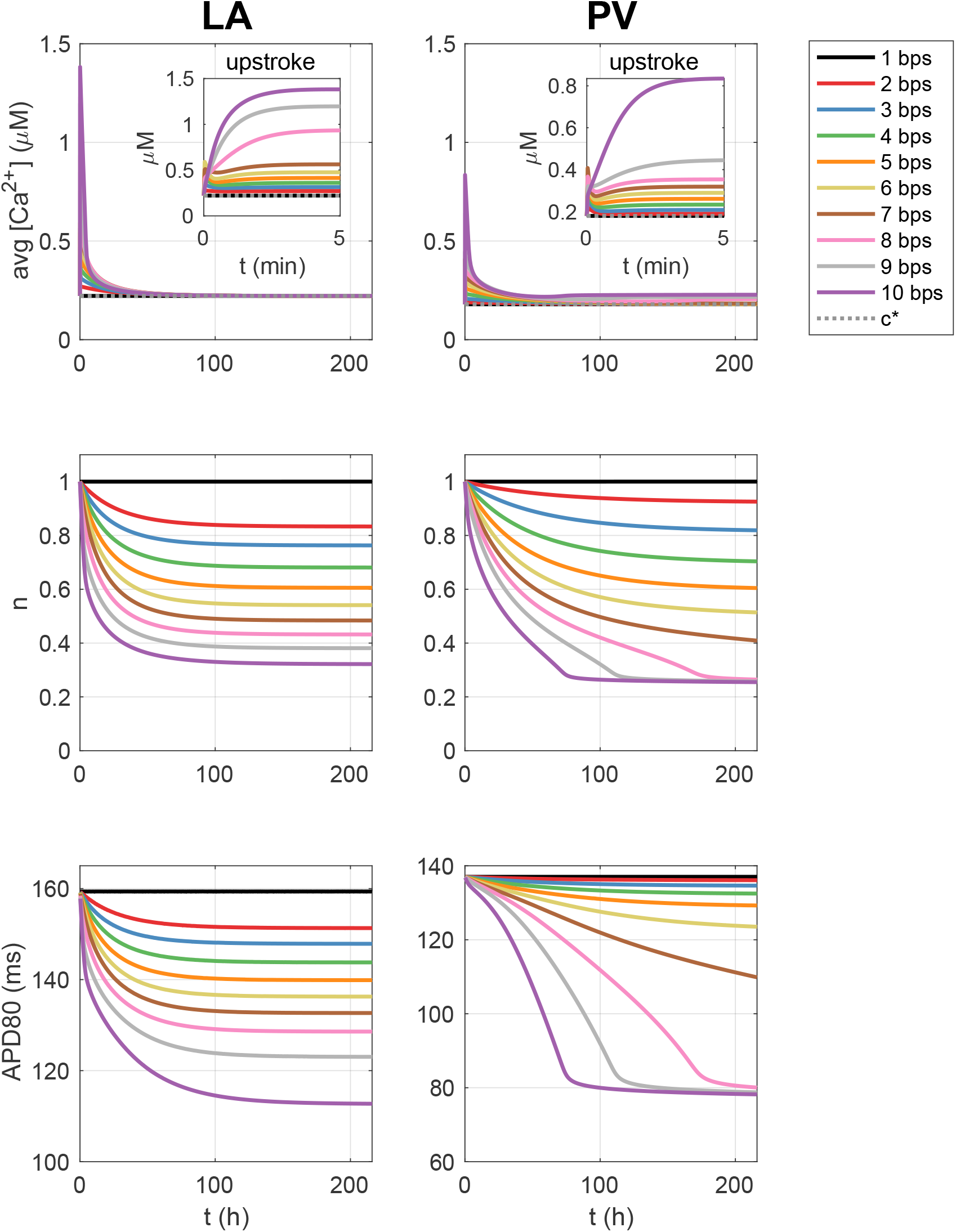
Membrane model simulations with rapid pacing. Effect of different pacing frequencies on the average (over one second) intracellular calcium concentration, the relative number of *I*_CaL_ channels, *n*, and the action potential duration at 80% repolarization (APD80) in membrane model simulations of the LA and PV models.

### 3.4 Increased re-entry susceptibility resulting from long-term rapid pacing

In Figure 7, we have plotted the membrane potential in the cells of the PV sleeve in SKNM simulations (see Figure 4B). The value of the relative number of *I*_CaL_ channels, *n*, for the PV and LA cells are collected from different points in time in membrane model simulations with pacing of 10 bps. We observe that after 2 h and 4 h of rapid pacing, the number of *I*_CaL_ channels is not reduced enough for a re-entrant wave to be generated, but after 6 h (and 8 h and 10 h) a re-entrant wave is generated.

**Figure 7:**
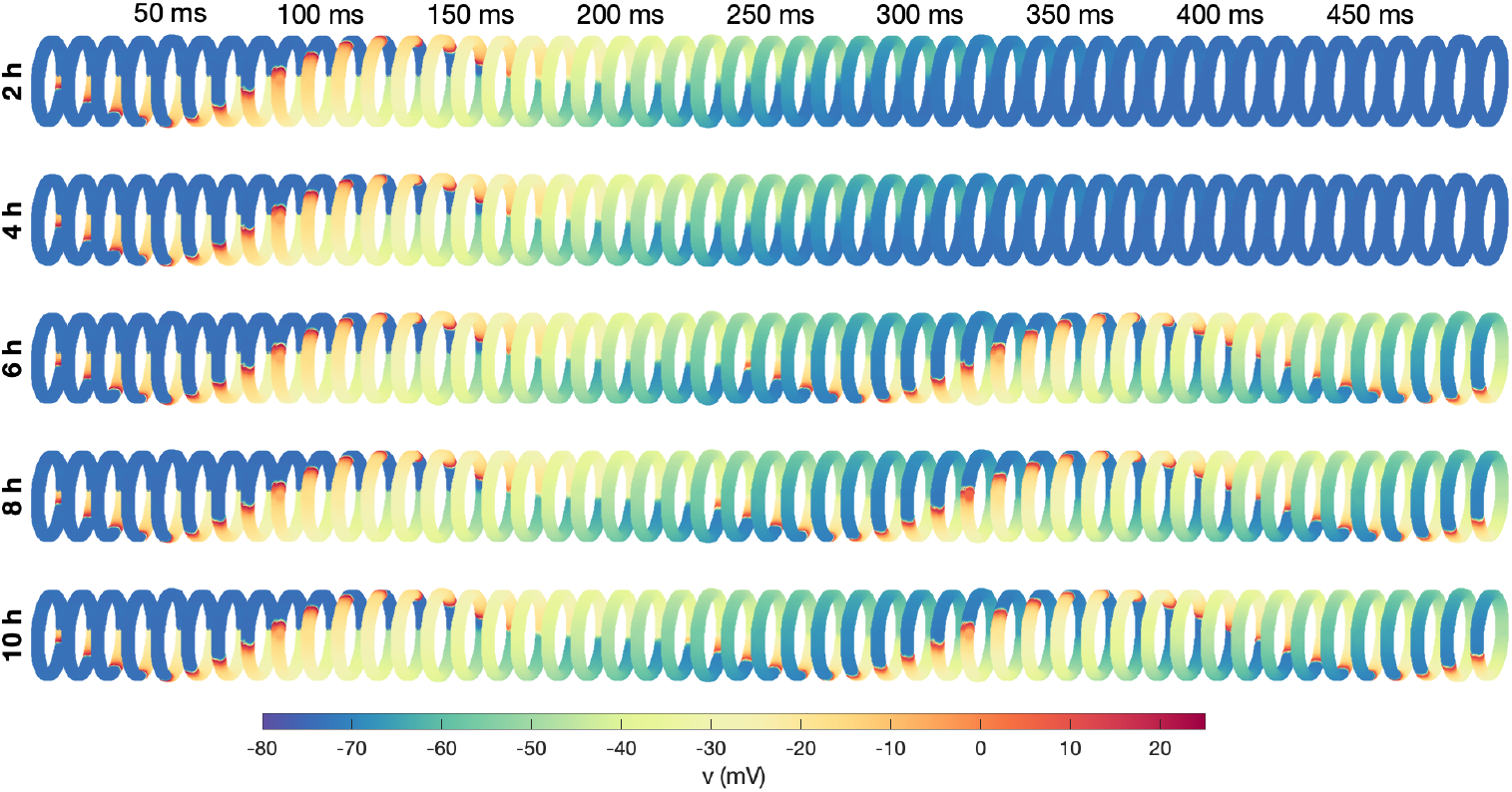
PV sleeve simulations after 2 h, 4 h, 6 h, 8 h and 10 h of rapid (10 bps) pacing. The figure shows the membrane potential for the cells in the PV sleeve cylinder at some different points in time after a stimulation is applied (indicated in the top text). Each row corresponds to a time point in the membrane model simulations of rapid pacing described in the text to the right. The simulation set-up is illustrated in Figure 4.

In Figure 8, we have performed the same type of simulation for a number of different time points and pacing frequencies and report whether or not a re-entrant wave is generated. We observe that the probability of re-entry increases as the number of beats per second increases and also when the duration of rapid pacing is increased.

**Figure 8:**
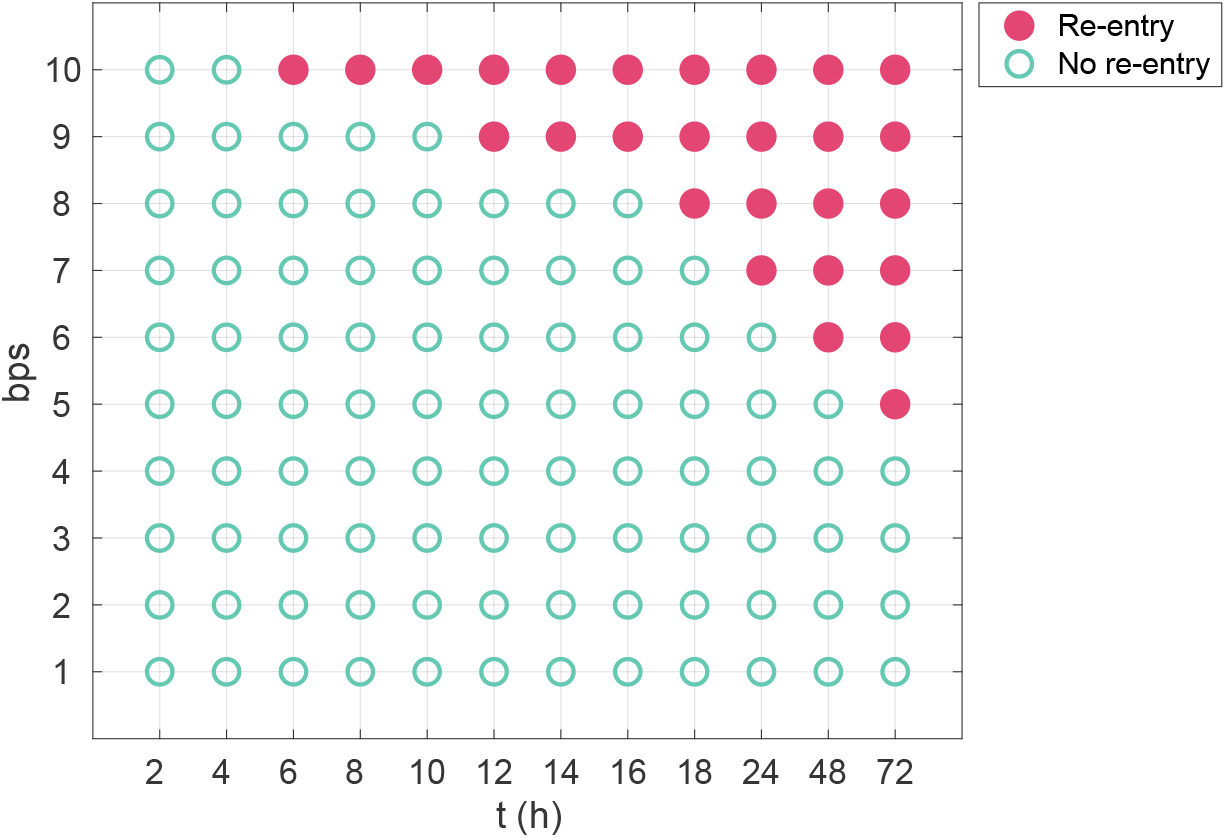
Re-entry susceptibility after hours of rapid pacing. The figure shows an overview of whether or not a re-entrant wave is generated after hours of different degrees of rapid pacing. Membrane model simulations are used to find the relative number of *I*_CaL_ channels, *n*, at the different points in time (see Figure 4A), and SKNM simulations of the PV sleeve is used to investigate whether a re-entrant wave is generated (see Figure 4B).

### 3.5 Influence of parameter perturbations

In Figure 9, we investigate how the results in Figure 8 depend on perturbations of essential model parameters.

**Figure 9:**
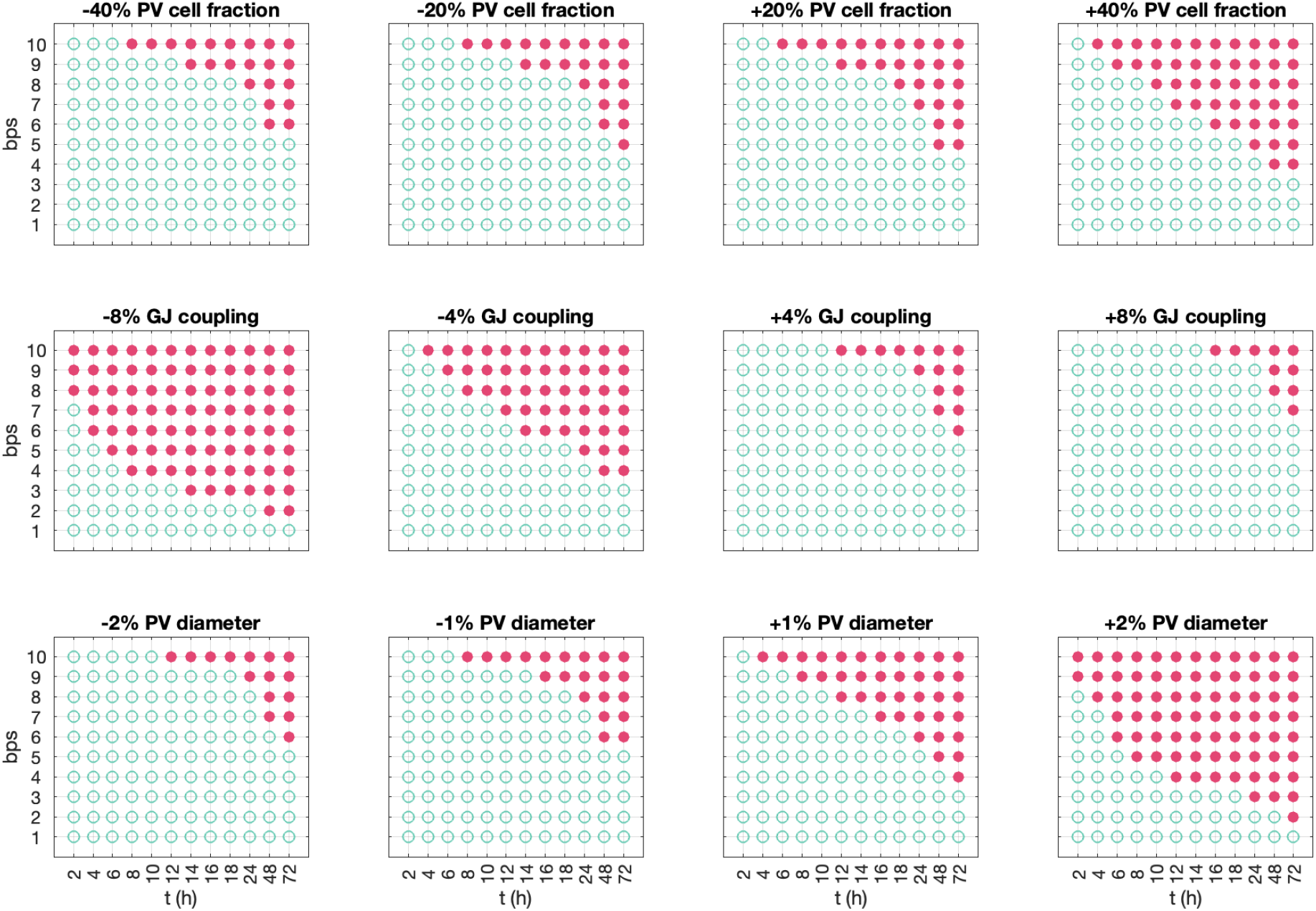
Re-entry susceptibility after hours of rapid pacing for variations of parameter values. The figure shows an overview of whether or not a re-entrant wave is generated after hours of different degrees of rapid pacing using the set-up described in Figure 4 for a number of adjusted parameter values. A closed red circle indicates that a re-entrant wave was generated, and an open green circle indicates that a re-entrant wave was not generated. In the upper panel, we have adjusted the fraction of PV myocytes in the PV sleeve from the default fraction of 50% PV myocytes and 50% LA myocytes. In the middle panel, we have adjusted the default strength of the gap junction coupling, 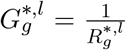 and 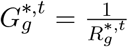, where 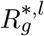 and 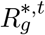 are given in Table 1. In the lower panel, we have varied the diameter of the PV sleeve by increasing or decreasing the number of surrounding cells.

#### Changing PV/LA-distribution

In the upper panel of Figure 9, we vary the fraction of PV myocytes in the PV sleeve, from the default fraction of 50% PV myocytes and 50% LA myocytes. We observe that as the fraction of PV myocytes increases, less time of rapid pacing and less frequent pacing is required before a re-entrant wave is generated.

#### Changing the gap junction density

Similarly, in the middle panel of Figure 9, we vary the strength of the default gap junction coupling between myocytes. We observe that if the gap junction coupling is, e.g., reduced by 8%, considerably less time of rapid pacing is required before to generate re-entry around the PV cylinder and considerably less frequent pacing is required. Conversely, if the gap junction coupling is increased by 8%, more frequent and long-lasting pacing is required to generate re-entry.

#### Changing the diameter of the pulmonary vein

Finally, in the lower panel of Figure 9, we vary the diameter of the PV sleeve by increasing of decreasing the number of surrounding myocytes. In this case, increasing the PV sleeve diameter considerably reduces the frequency and duration of rapid pacing required to induce a re-entrant wave around the PV sleeve.

## 4 Discussion

It is well-established that atrial fibrillation (AF) follows the concept of *AF begets AF*, signifying its progressive and self-perpetuating nature, see, e.g., [54, 55, 56, 57]. This implies that episodes of AF contribute to the development of a conducive substrate for subsequent occurrences, thereby rendering the condition progressively more entrenched and potentially leading to a self-sustaining mechanism. Here, we have proposed a pathway from paroxysmal AF episodes to sustained re-entry at the left pulmonary vein’s outlet, employing a mathematical model to expose this progression, see Figure 1.

### Reduced *I*_CaL_ can induce re-entry

In Figure 3, we show how excitation waves can propagate around the pulmonary vein outlet. We examine two scenarios: a control case and one where the calcium current, *I*_CaL_, is halved. A reduction in *I*_CaL_ is associated with a decrease in action potential duration (APD), which, as demonstrated in the simulation depicted in Figure 3, leads to a heightened risk of re-entry wave formation. The relationship between diminished calcium current and shortened APD has been well-documented, see [58, 59]. This, in turn, results in a reduced effective refractory period, escalating the likelihood of re-entrant wave emergence.

### Rapid firing reduce the calcium current

The effect of rapid firing on the strength of the calcium current as modelled by (10) is shown in Figure 5. Rapid firing leads to increased cytosolic calcium load which according to the model (10) reduces the calcium current. The effect of increased cytosolic calcium load during rapid firing is well documented, [59], and easily seen in simulations based on AP models of atrial cells, see Figure S3 in the Supplementary Information. Weakened calcium current during rapid firing is also well documented, see, e.g., [17, 60, 59, 18, 61]. Here we suggest that the remodeling can be represented by (10). According to this model, the calcium current, *I*_CaL_, is down-regulated when the cytosolic calcium concentration is elevated above its target value.

### Stable re-entry after hours of rapid firing

In Figure 7, we demonstrate that re-entry may develop after extended periods of rapid firing. For this example, we simulated the membrane model (no spatial variation) continuously for up to ten hours to monitor the impact of sustained rapid firing on the calcium current dynamics. At intervals of 2 h, 4 h, 6 h, 8 h, and 10 h, we analyzed the spatial model’s behavior. It was noted that at the 2-hour and 4-hour marks, the excitation waves ceased after a single circumnavigation of the pulmonary vein. However, with prolonged rapid firing – specifically after 6 h, 8 h, and 10 h – a consistent re-entry pattern emerged. This pattern suggests that the cellular remodeling induced by the rapid firing persists for several hours before the re-entry stabilizes. In Figure 8, we show how the path to re-entry depends on the frequency and duration of the firing. More prolonged, and faster, rapid firing clearly increases the occurrence of re-entry.

### The tissue properties strongly influence the occurrence of re-entry

Modifying the properties of the tissue significantly alters the required duration and frequency of rapid firing to initiate re-entry. Figure 9 shows that the ratio of pulmonary vein (PV) to left atrium (LA) myocytes has a decisive impact on the propensity for re-entry. An increased proportion of PV myocytes is associated with a higher incidence of re-entry events. Similarly, a reduction in gap junction coupling correlates with a more frequent occurrence of re-entry. The diameter of the pulmonary vein is particularly critical; a slight enlargement markedly elevates the incidence of re-entry.

### Remodeling of other currents

In the models presented in [24] and [31], regulation by cytosolic calcium concentration extends beyond the calcium current. In [24] regulation of all membrane currents was included, while [31] in addition allowed aspects of intracellular calcium dynamics to be controlled by the calcium concentration. Although our primary focus remains on the calcium current, as in [32], we acknowledge that other currents and fluxes may also be impacted. In the Supplementary Information (Figure S4) we show the effect of increasing the individual membrane currents by 20%. As seen in that figure, increase in the *I*_CaL_ current, increases the intracellular calcium concentration. Therefore, regulation of calcium overload by decreasing the number of calcium channels is reasonable. The same effect holds for the background calcium current. But the fast sodium current *I*_Na_ in both LA and PV cardiomyocytes have very limited effects on the intracellular calcium concentrations. Therefore, regulation of this current in order to repair deviation from the calcium target value appears to be ineffectual. Several of the potassium currents work the other way around compared to *I*_CaL_ and cannot be used to control intracellular calcium in the same way as the calcium current. A comprehensive review of cardiac ion channel remodeling is given in [61] (see Table 5). For atrial cardiomyocytes, *I*_CaL_ is weakened during AF, but the potassium currents have more complex behavior. For instance, the *I*_to_ current (transient outward potassium current) is decreased, but the *I*_K1_ current (inward rectifier potassium current) is increased. To conclude, it is not straightforward to use the model (10) for all the currents of the atrial cell membrane in the way suggested by [24] and [31], and we therefore restrict our attention to the *I*_CaL_ current. These considerations and the conclusion are in line with the discussion of similar issues in [32].

## 5 Conclusion

We have presented a computational model that demonstrates a possible path from paroxysmal AF episodes to sustained atrial fibrillation. The path appears to be robust in the sense that re-entry is obtained under a series of reasonable perturbations to the parameters. The model predicts that the occurrence of re-entry is proportional to the frequency and duration of the rapid firing. The occurrence also increases when the gap junction coupling is decreased and when the diameter of the pulmonary vein is increased. Finally, it increases as the fraction of pulmonary vein myocytes in relation to left atrial myocytes increases. The progression of atrial fibrillation, as depicted by the vicious cycle in Figure 1, underscores the critical importance of early intervention in this cycle. The likelihood of successful repair diminishes as the disease advances, highlighting the necessity of timely therapeutic measures.

## Supplementary Information

### S1 Model formulation

In this supplementary section, we describe the formulation of the human left atrial (LA) and pulmonary vein (PV) cardiomyocyte versions of our base model. Here, the membrane potential (*v*) is given in units of mV, and the Ca^2+^ and Na^+^ concentrations are given in units of mM. All currents are given in units of A/F, and the ionic fluxes are expressed as mmol/ms per total cell volume (i.e., in units of mM/ms). Time is given in ms. The parameters of the model are all given in Tables S1–S7. In particular, the adjustment factors used to scale the model from the left atrial version to the pulmonary vein version of the model are found in Table S4.

Note that the model formulation is very similar to the base model from [33]. The only difference is that *Ī*_NaCa_ is decreased by 40%, *g*_Kr_ is decreased by 20% and the calcium buffer concentrations 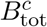 and 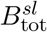 are increased by 40% compared to the version in [33]. In addition, a model for the number of *I*_CaL_ channels on the cell membrane has been added.

#### S1.1 Membrane potential

In the base model formulation, the membrane potential is governed by

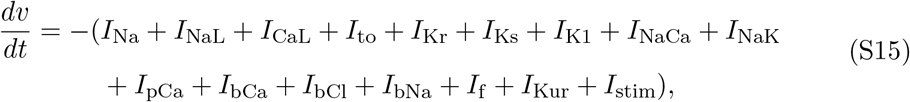

where *I*_Na_, *I*_NaL_, *I*_CaL_, *I*_to_, *I*_Kr_, *I*_Ks_, *I*_K1_, *I*_NaCa_, *I*_NaK_, *I*_pCa_, *I*_bCl_, *I*_bCa_, *I*_bNa_, *I*_f_, and *I*_Kur_ are transmembrane currents that will be specified below and *I*_stim_ is an applied stimulus current. Unless otherwise specified, we let *I*_stim_ be given as a constant current of size −40 A/F applied for 1 ms.

#### S1.2 Transmembrane currents

In general, the currents through the voltage-gated ion channels within the myocyte membrane are given on the form

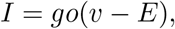

where *g* is the channel conductance, *v* is the membrane potential and *E* is the equilibrium potential of the channel. Moreover, *o* is the open probability of the channels, which is given on the form *o* = Π _*i*_ *z*_*i*_, where *z*_*i*_ are gating variables. These gating variables are either given as an explicit function of the membrane potential or governed by equations of the form

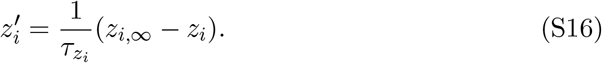

The parameters 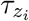 and *z*_*i*,∞_ are specified for each of the gating variables of the model in Table S10.

##### Fast sodium current (*I*_Na_)

The formulation of the fast sodium current is based on the model formulation given in [62] and is given by

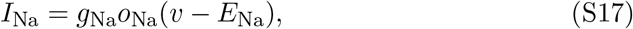

where the open probability is given by

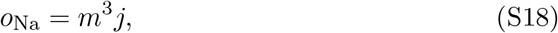

and *m* and *j* are gating variables governed by equations of the form (S16).

##### Late sodium current (*I*_NaL_)

The formulation of the late sodium current, *I*_NaL_, is based on [63] and is given by

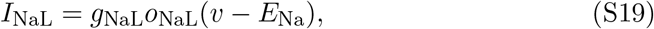

where the open probability is given by

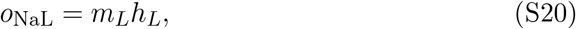

and *m*_*L*_ and *h*_*L*_ are gating variables governed by equations of the form (S16).

##### Transient outward potassium current (*I*_to_)

The formulation of the transient outward potassium current, *I*_to_, is based on [64] and is given by

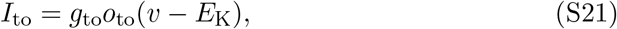

where the open probability is given by

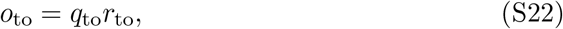

and *q*_to_ and *r*_to_ are gating variables governed by equations of the form (S16).

##### Rapidly activating potassium current (*I*_Kr_)

The rapidly activating potassium current, *I*_Kr_, is formulated as a Markov model, based on [65]. The formulation has been fitted to data of WT and N588K *I*_Kr_ currents from [66]. The current is given by

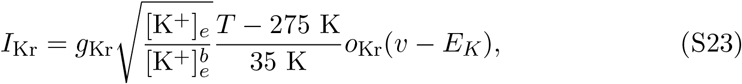

where *o*_Kr_ is modelled by a Markov model of the form

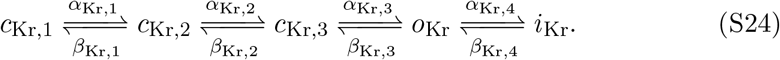

Here, the dynamics of the closed states *c*_Kr,1_, *c*_Kr,2_, and *c*_Kr,3_, the open state *o*_Kr_, and the inactivated state *i*_Kr_ are given by

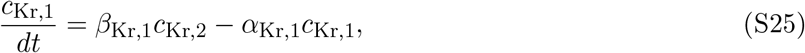

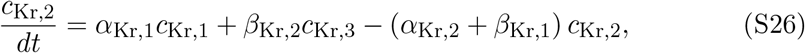

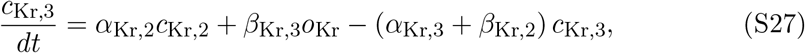

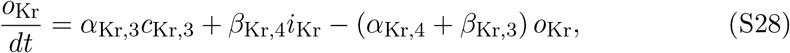

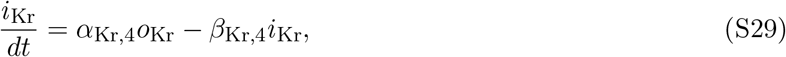

where the transition rates are given by provided in Table S8.

##### Slowly activating potassium current (*I*_Ks_)

The formulation of the slowly activating potassium current, *I*_Ks_, is based on [62] and is given by

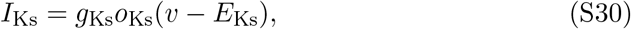

where

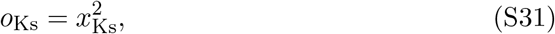

and the dynamics of *x*_Ks_ is governed by an equation of the form (S16).

##### Inward rectifier potassium current (*I*_K1_)

The formulation of the inward rectifier potassium current, *I*_K1_, is based on [67, 68], and is given by

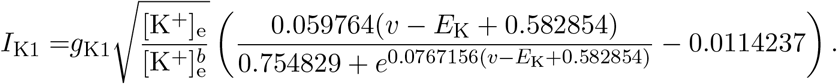

##### Ultrarapid delayed rectifier potassium current (*I*_Kur_)

The formulation of the ultrarapid delayed rectifier potassium current, *I*_Kur_, is based on [69, 70] and is given by

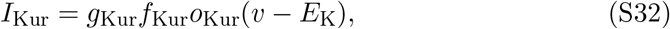

where *f*_Kur_ is given by

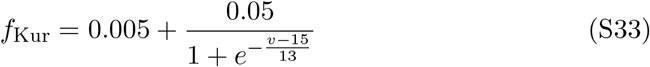

and

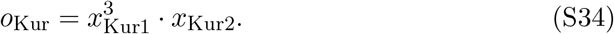

The dynamics of *x*_Kur1_ and *x*_Kur2_ are governed by equations of the form (S16).

##### Hyperpolarization activated funny current (*I*_f_)

The formulation for the hyperpolarization activated funny current, *I*_f_, is based on [64] and is given by

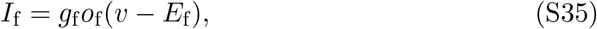

where

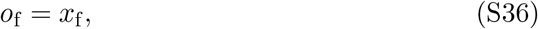

and the dynamics of *x*_f_ is governed by an equation of the form (S16).

##### L-type Ca^2+^ current (*I*_CaL_)

The formualtion for the L-type Ca^2+^ current, *I*_CaL_, is based on the formulation in [62] and is given by

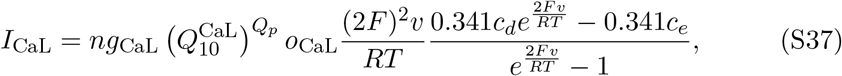

where

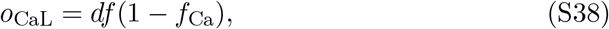

and the dynamics of *d, f* and *f*_Ca_ are governed by equations of the form (S16). Note that *n* is governed by (S68) and represents the relative number of *I*_CaL_ channels.

##### Time-independent background currents (*I*_bCa_, *I*_bBa_, *I*_bCl_)

The formulation of the background currents, *I*_bCa_, *I*_bNa_ and *I*_bCl_, are based on [62] and are given by

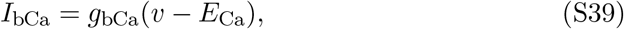

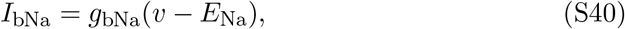

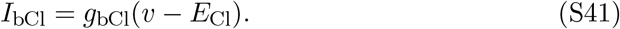

##### Sodium-calcium exchanger current (*I*_NaCa_)

The formulation of the Na^+^-Ca^2+^ exchanger current, *I*_NaCa_, is based on [62] and is given by

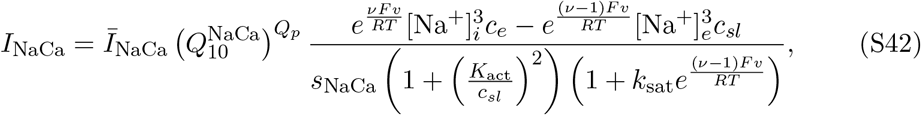

where

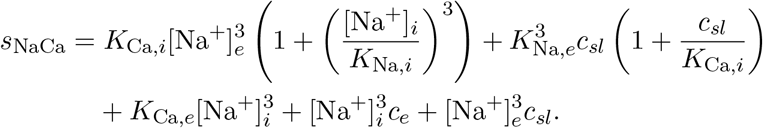

##### Sarcolemmal Ca^2+^ pump current (*I*_pCa_)

The formulation of the current through the sarcolemmal Ca^2+^ pump, *I*_pCa_, is based on [62] and is given by

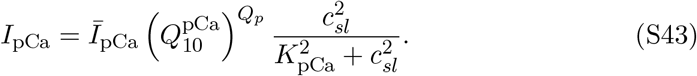

##### Sodium-potassium pump current (*I*_NaK_)

The current through the Na^+^-K^+^ pump, *I*_NaK_, is based on [62] and is given by

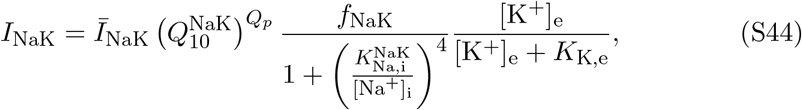

where

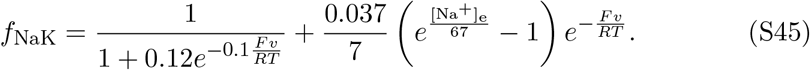

#### S1.3 Intracellular [Ca^2+^] dynamics

The Ca^2+^ dynamics are governed by

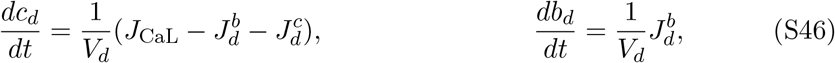

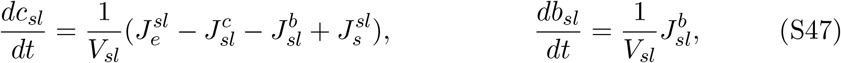

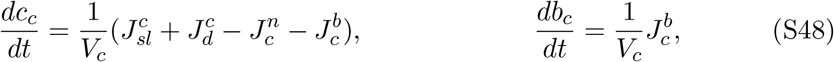

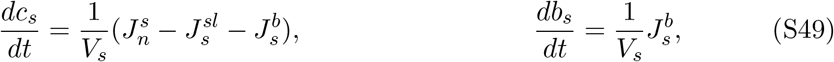

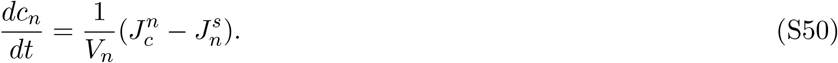

Here, *c*_*d*_ is the concentration of free Ca^2+^ in the dyad, *b*_*d*_ is the concentration of Ca^2+^ bound to a buffer in the dyad, *c*_*sl*_ is the concentration of free Ca^2+^ in the sub-sarcolemmal (SL) compartment, *b*_*sl*_ is the concentration of Ca^2+^ bound to a buffer in the SL compartment, *c*_*c*_ is the concentration of free Ca^2+^ in the bulk cytosol, *b*_*c*_ is the concentration of Ca^2+^ bound to a buffer in the bulk cytosol, *c*_*s*_ is the concentration of free Ca^2+^ in the junctional sarcoplasmic reticulum (jSR), *b*_*s*_ is the concentration of Ca^2+^ bound to a buffer in the jSR, and *c*_*n*_ is the concentration of free Ca^2+^ in the network sarcoplasmic reticulum (nSR). The expressions for the fluxes are specified below.

#### S1.4 Ca^2+^ fluxes

##### Flux through the SERCA pumps

The flux from the bulk cytosol to the nSR through the SERCA pumps is based on [62] and given by

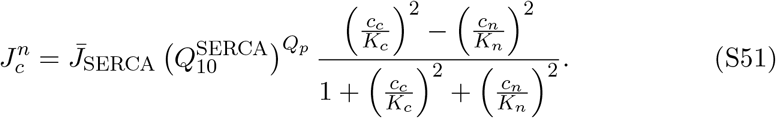

##### Flux through the RyRs

The flux from the jSR to the SL compartment is given by

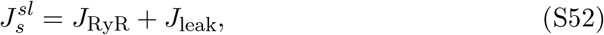

where *J*_RyR_ is the flux through the active RyR channels and *J*_leak_ is the flux through passive RyR channels that are always open, given by

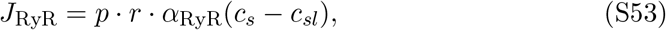

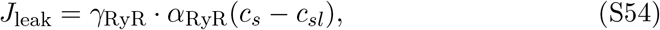

respectively. Here, *p* represents the open probability of the active RyR channels and is given by

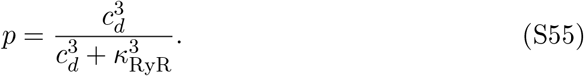

Furthermore, *r* is the fraction of RyR channels that are not inactivated and is governed by the equation

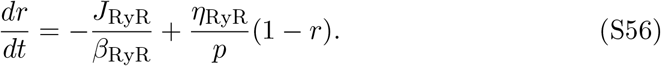

##### Passive diffusion fluxes between compartments

The passive diffusion fluxes between intracellular compartments are given by

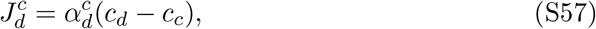

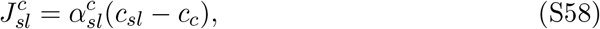

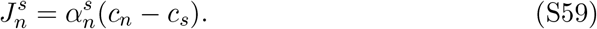

##### Buffer fluxes

The fluxes of free Ca^2+^ binding to a Ca^2+^ buffer are given by

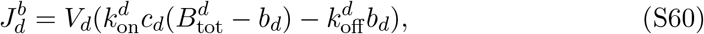

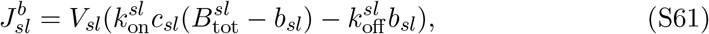

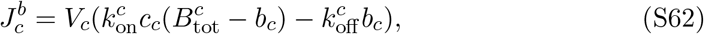

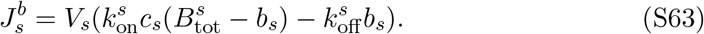

**Figure S1:**
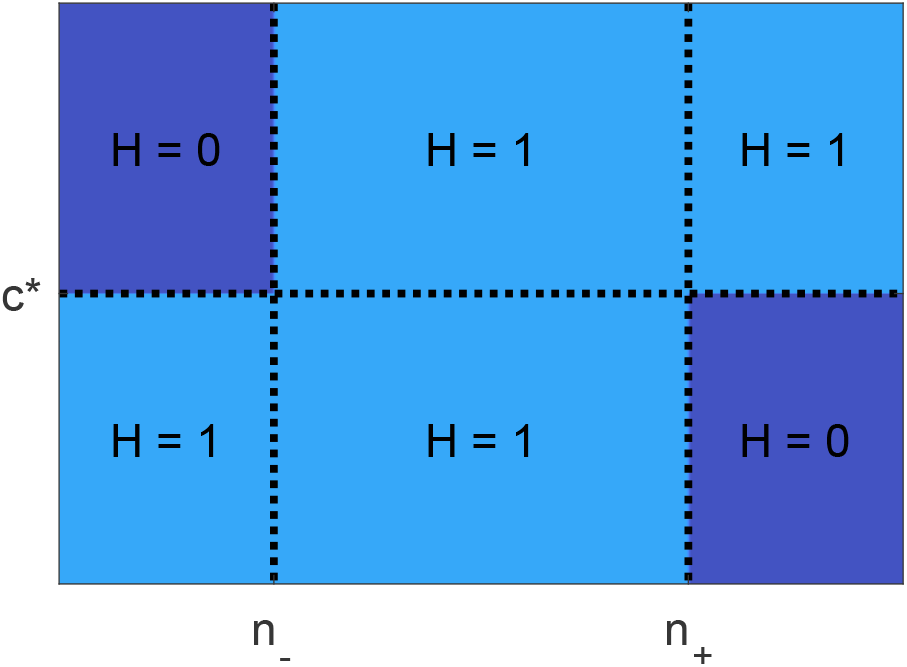
Illustration of the function *H*(*c, n*).

##### Membrane fluxes

The membrane Ca^2+^ fluxes, *J*_CaL_, *J*_bCa_, *J*_pCa_, and *J*_NaCa_, are given by

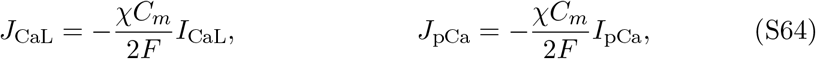

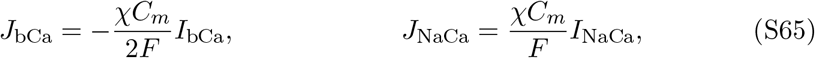

where *I*_CaL_, *I*_bCa_, *I*_pCa_, and *I*_NaCa_ are defined by the expressions given above. Furthermore,

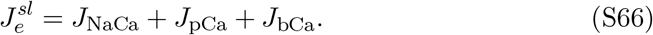

#### S1.5 Intracellular Na^+^ dynamics

The intracellular Na^+^ concentration is governed by

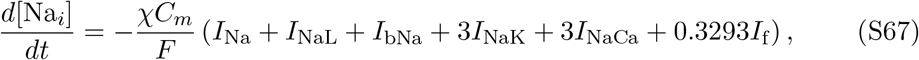

where the currents *I*_Na_, *I*_NaL_, *I*_bNa_, *I*_NaK_, *I*_NaCa_, and *I*_f_ are specified above.

#### S1.6 Protein regulation

The scaling factor *n* for the number of L-type calcium channels on the cell membrane is modeled by

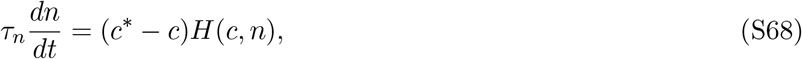

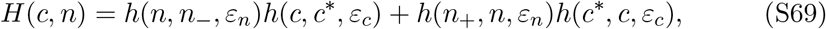

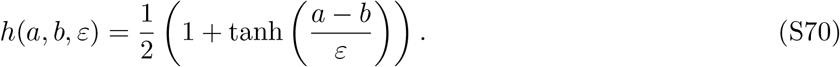

The function *H*(*c, n*) is illustrated in Figure S1.

#### S1.7 Nernst equilibrium potentials

The Nernst equilibrium potentials for the ion channels are defined as

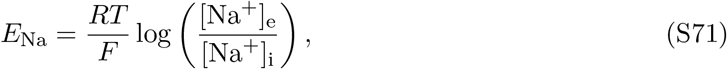

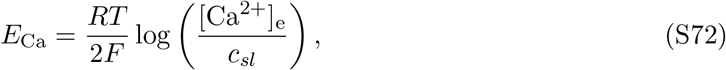

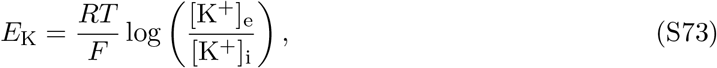

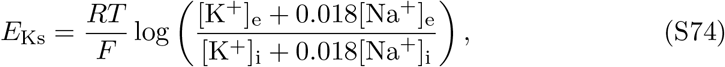

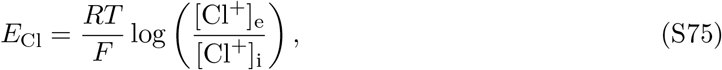

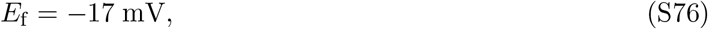

for the parameter values given in Table S2.

#### S1.8 Baseline parameter values

**Table S1:** Default geometry parameters of the base model.

| Parameter | Description | Value |
| --- | --- | --- |
| $V_d$ | Volume fraction of the dyadic subspace | 0.001 |
| $V_{sl}$ | Volume fraction of the SL compartment | 0.028 |
| $V_c$ | Volume fraction of the bulk cytosol | 0.917 |
| $V_s$ | Volume fraction of the jSR | 0.004 |
| $V_n$ | Volume fraction of the nSR | 0.05 |
| $\chi$ | Cell surface to volume ratio | $0.6 \mu\text{m}^{-1}$ |

### S2 Comparison of KNM and SKNM solutions

Figure S2 shows a comparison of the solutions of the Kirchhoff network model (KNM) and the simplified Kirchhoff network model (SKNM) [41, 34].

### S3 Properties of the LA and PV membrane models with a fixed number of channels

In this section, we investigate some properties of the LA and PV membrane models described in Section S1. We consider the version of the models without protein regulation (i.e., when the number of calcium channels is fixed at *n* = 1).

**Table S2:** Physical constants and ionic concentrations of the base model.

| Parameter | Description | Value |
| --- | --- | --- |
| $C_m$ | Specific membrane capacitance | 0.01 pF/$\mu\text{m}^2$ |
| $F$ | Faraday's constant | 96.485 C/mmol |
| $R$ | Universal gas constant | 8.314 J/(mol·K) |
| $T$ | Temperature | 310 K |
| $[\text{Ca}^{2+}]_e$ | Extracellular $\text{Ca}^{2+}$ concentration | 1.8 mM |
| $[\text{Na}^+]_e$ | Extracellular sodium concentration | 140 mM |
| $[\text{K}^+]_e$ | Extracellular potassium concentration | 5.4 mM |
| $[\text{K}^+]_e^b$ | Baseline extracellular potassium concentration | 5.4 mM |
| $[\text{K}^+]_i$ | Intracellular potassium concentration | 120 mM |
| $[\text{Cl}^-]_e$ | Extracellular chloride concentration | 150 mM |
| $[\text{Cl}^-]_i$ | Intracellular chloride concentration | 15 mM |

**Table S3:** Conductances and similar cell-specific parameter values in the base model formulation. Note that the parameter values of this table define the human left atrial version of the base model. For the human pulmonary vein version, the adjustment factors of Table S4 are applied.

| Parameter | Value | Parameter | Value |
| --- | --- | --- | --- |
| $g_{\text{Na}}$ | 7.056 mS/$\mu\text{F}$ | $\bar{I}_{\text{NaCa}}$ | 2.22 $\mu\text{A}/\mu\text{F}$ |
| $g_{\text{NaL}}$ | 0.021 mS/$\mu\text{F}$ | $\bar{I}_{\text{pCa}}$ | 0.068 $\mu\text{A}/\mu\text{F}$ |
| $g_{\text{to}}$ | 0.55 mS/$\mu\text{F}$ | $\bar{J}_{\text{SERCA}}$ | 0.00024 mM/ms |
| $g_{\text{Kr}}$ | 1.2 mS/$\mu\text{F}$ | $\alpha_{\text{RyR}}$ | 0.0075 $\text{ms}^{-1}$ |
| $g_{\text{Ks}}$ | 0.21 mS/$\mu\text{F}$ | $\beta_{\text{RyR}}$ | 0.038 mM |
| $g_{\text{K1}}$ | 2.62 mS/$\mu\text{F}$ | $\alpha_d^c$ | 0.0034 $\text{ms}^{-1}$ |
| $g_{\text{f}}$ | 0.0001 mS/$\mu\text{F}$ | $\alpha_{sl}^c$ | 0.15 $\text{ms}^{-1}$ |
| $g_{\text{bCl}}$ | 0.0114 mS/$\mu\text{F}$ | $\alpha_n^s$ | 0.012 $\text{ms}^{-1}$ |
| $g_{\text{CaL}}$ | 0.049 nL/($\mu\text{F ms}$) | $B_{\text{tot}}^c$ | 0.07 mM |
| $g_{\text{bCa}}$ | 0.000832 mS/$\mu\text{F}$ | $B_{\text{tot}}^d$ | 1.2 mM |
| $g_{\text{bNa}}$ | 0.00185 mS/$\mu\text{F}$ | $B_{\text{tot}}^{sl}$ | 0.9 mM |
| $g_{\text{Kur}}$ | 2.475 mS/$\mu\text{F}$ | $B_{\text{tot}}^s$ | 27 mM |
| $\bar{I}_{\text{NaK}}$ | 2.76 $\mu\text{A}/\mu\text{F}$ | | |

**Table S4:** Adjustment factors for the pulmonary vein (PV) version of the base model, based on [44].

| Parameter | PV scaling factor |
| --- | --- |
| $g_{K1}$ | 0.58 |
| $g_{Kr}$ | 1.5 |
| $g_{Ks}$ | 1.6 |
| $g_{to}$ | 0.75 |
| $g_{CaL}$ | 0.7 |

**Table S5:** Parameters for the intracellular Ca^2+^ fluxes of the base model.

| Parameter | Flux | Value |
| --- | --- | --- |
| $K_c$ | $J_c^n$ | 0.00025 mM |
| $K_n$ | $J_c^n$ | 1.7 mM |
| $\gamma_{RyR}$ | $J_s^{sl}$ | 0.001 |
| $\kappa_{RyR}$ | $J_{RyR}$ | 0.015 mM |
| $\eta_{RyR}$ | $J_s^{sl}$ | 0.00001 ms<sup>-1</sup> |

**Table S6:** Additional parameters for the membrane currents of the base model.

| Parameter | Current | Value |
| --- | --- | --- |
| $k_{sat}$ | $I_{NaCa}$ | 0.3 |
| $\nu$ | $I_{NaCa}$ | 0.3 |
| $K_{act}$ | $I_{NaCa}$ | 0.00015 mM |
| $K_{Ca,i}$ | $I_{NaCa}$ | 0.0036 mM |
| $K_{Ca,e}$ | $I_{NaCa}$ | 1.3 mM |
| $K_{Na,i}$ | $I_{NaCa}$ | 12.3 mM |
| $K_{Na,e}$ | $I_{NaCa}$ | 87.5 mM |
| $K_{Na,i}^{NaK}$ | $I_{NaK}$ | (11 mM) · ($Q_{10}^{KNaK}$)<sup>$Q^p$</sup> |
| $K_{K,e}$ | $I_{NaK}$ | 1.5 mM |
| $K_{pCa}$ | $I_{pCa}$ | 0.0005 mM |

**Table S7:** Transition rates for the Ca^2+^ buffers of the base model.

| Parameter | Compartment | Value |
| --- | --- | --- |
| $k_{\text{on}}^c$ | Bulk cytosol | $40 \text{ ms}^{-1}\text{mM}^{-1}$ |
| $k_{\text{off}}^c$ | Bulk cytosol | $0.03 \text{ ms}^{-1}$ |
| $k_{\text{on}}^d$ | Dyad | $100 \text{ ms}^{-1}\text{mM}^{-1}$ |
| $k_{\text{off}}^d$ | Dyad | $1 \text{ ms}^{-1}$ |
| $k_{\text{on}}^{sl}$ | Subsarcolemmal space | $100 \text{ ms}^{-1}\text{mM}^{-1}$ |
| $k_{\text{off}}^{sl}$ | Subsarcolemmal space | $0.15 \text{ ms}^{-1}$ |
| $k_{\text{on}}^s$ | Junctional SR | $100 \text{ ms}^{-1}\text{mM}^{-1}$ |
| $k_{\text{off}}^s$ | Junctional SR | $65 \text{ ms}^{-1}$ |

**Table S8:** Transition rates for the *I*_Kr_ Markov model. Here,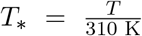, where *T* is the temperature.

|  |  |
| --- | --- |
| $\alpha_{\text{Kr},1}$ | $0.4 \cdot T_* \cdot e^{24.335+T_*^{-1}(0.0112v-25.914)}$ |
| $\beta_{\text{Kr},1}$ | $T_* \cdot e^{13.688+T_*^{-1}(-0.0603v-15.707)}$ |
| $\alpha_{\text{Kr},2}$ | $0.4 \cdot T_* \cdot e^{22.746+T_*^{-1}(-25.914)}$ |
| $\beta_{\text{Kr},2}$ | $T_* \cdot e^{13.193+T_*^{-1}(-15.707)}$ |
| $\alpha_{\text{Kr},3}$ | $0.4 \cdot T_* \cdot e^{22.098+T_*^{-1}(0.0365v-25.914)}$ |
| $\beta_{\text{Kr},3}$ | $T_* \cdot e^{7.313+T_*^{-1}(-0.0399v-15.707)}$ |
| $\alpha_{\text{Kr},4}$ | $T_* \cdot \left( \frac{[\text{K}^+]_e^b}{[\text{K}^+]_e} \right)^{0.4} e^{30.016+T_*^{-1}(0.0223v-30.888)}$ |
| $\beta_{\text{Kr},4}$ | $T_* \cdot e^{30.061+T_*^{-1}(-0.0312v-33.243)}$ |

**Table S9:** Parameters for the regulation of the number of L-type calcium channels

| Parameter | Value | Parameter | Value |
| --- | --- | --- | --- |
| $n_-$ | 0.28 | $n_+$ | 3 |
| $\varepsilon_n$ | 0.01 | $\varepsilon_c$ | $10^{-7} \text{ mM}$ |
| $c^*$ (LA) | $2.22 \cdot 10^{-4} \text{ mM}$ | $c^*$ (PV) | $1.79 \cdot 10^{-4} \text{ mM}$ |
| $\tau_n$ | 30,000 mMms | | |

**Table S10:** Specification of the parameters *z*_∞_ and *τ*_*z*_, for *z* = *m, j, m*_*L*_, *h*_*L*_, *d, f, f*_Ca_, *q*_to_, *r*_to_, *x*_Ks_, *x*_Kur1_, *x*_Kur2_, and *x*_f_ in the equations for the gating variables (S16).

| Current | Gate | $z_\infty$ | $\alpha_z$ | $\beta_z$ | $\tau_z$ |
| --- | --- | --- | --- | --- | --- |
| $I_{\text{Na}}$ | $m$ | $\frac{1}{(1 + e^{(-57-v)/9})^2}$ | $0.13e^{-((v+46)/16)^2}$ | $0.06e^{-((v-5)/51)^2}$ | $\frac{\alpha_m + \beta_m}{(Q_{10}^{\text{Na}})^{Q_p}}$ |
| | $j$ | $\frac{1}{(1 + e^{(v+72)/7})^2}$ | $\begin{cases} 0, & \text{if } v \geq -40 \\ \frac{(-2.5 \cdot 10^4 e^{0.2v} - 7 \cdot 10^{-6} e^{-0.04v})(v+38)}{1 + e^{0.3(v+79)}}, & \text{otherwise} \end{cases}$ | $\begin{cases} \frac{0.6e^{0.06v}}{1 + e^{-0.1(v+32)}}, & \text{if } v \geq -40 \\ \frac{0.02e^{-0.01v}}{1 + e^{-0.14(v+40)}}, & \text{otherwise} \end{cases}$ | $\frac{1}{(\alpha_j + \beta_j)(Q_{10}^{\text{Na}})^{Q_p}}$ |
| $I_{\text{NaL}}$ | $m_L$ | $\frac{1}{1 + e^{(-43-v)/5}}$ | $\frac{1}{6.8e^{(v+12)/35}}$ | $8.6e^{-(v+77)/6}$ | $\frac{\alpha_m + \beta_m}{(Q_{10}^{\text{NaL}})^{Q_p}}$ |
| | $h_L$ | $\frac{1}{1 + e^{(v+88)/7.5}}$ | | | $\frac{200 \text{ ms}}{(Q_{10}^{\text{NaL}})^{Q_p}}$ |
| $I_{\text{CaL}}$ | $d$ | $\frac{1}{1 + e^{-(v+20)/6}}$ | $\frac{1 - e^{-\frac{v+5}{6}}}{0.035(v+5)}$ | | $\alpha_d d_\infty$ |
| | $f$ | $\frac{1}{1 + e^{(v+35)/9}} + \frac{0.6}{1 + e^{(50-v)/20}}$ | $\frac{1}{0.02e^{-(0.034(v+14.5)^2)} + 0.02}$ | | $\alpha_f$ |
| | $f_{\text{Ca}}$ | $\frac{1.7c_d}{1.7c_d + 0.012}$ | $\frac{1}{1.7c_d^{1.5} + 0.012}$ | | $\alpha_{\text{Ca}}$ |
| $I_{\text{to}}$ | $q_{\text{to}}$ | $\frac{1}{1 + e^{(v+53)/13}}$ | $\frac{39}{0.57e^{-0.08(v+44)} + 0.065e^{0.1(v+46)}}$ | 6 | $\frac{\alpha_{q_{\text{to}}} + \beta_{q_{\text{to}}}}{(Q_{10}^{\text{to}})^{Q_p}}$ |
| | $r_{\text{to}}$ | $\frac{1}{1 + e^{-(v-22.3)/18.75}}$ | $\frac{14.4}{e^{0.09(v+30.61)} + 0.37e^{-0.12(v+24)}}$ | 2.75 | $\frac{\alpha_{r_{\text{to}}} + \beta_{r_{\text{to}}}}{(Q_{10}^{\text{to}})^{Q_p}}$ |
| $I_{\text{Ks}}$ | $x_{\text{Ks}}$ | $\frac{1}{1 + e^{-(v+3.8)/14}}$ | $\frac{990}{1 + e^{-(v+2.4)/14}}$ | | $\frac{\alpha_{x_{\text{Ks}}}}{(Q_{10}^{\text{Ks}})^{Q_p}}$ |
| $I_{\text{Kur}}$ | $x_{\text{Kur1}}$ | $\frac{1}{1 + e^{-(v+30.3)/9.6}}$ | $\frac{0.65}{e^{-(v+10)/8.5} + e^{-(v-30)/59}}$ | $\frac{0.65}{2.5 + e^{(v+82)/17}}$ | $\frac{1}{(\alpha_{x_{\text{Kur1}}} + \beta_{x_{\text{Kur1}}})(Q_{10}^{\text{Kur}})^{Q_p}}$ |
| | $x_{\text{Kur2}}$ | $\frac{1}{1 + e^{(v-99.45)/27.48}}$ | $\frac{1}{21 + e^{-(v-185)/28}}$ | $e^{(v-158)/16}$ | $\frac{1}{(\alpha_{x_{\text{Kur2}}} + \beta_{x_{\text{Kur2}}})(Q_{10}^{\text{Kur}})^{Q_p}}$ |
| $I_{\text{f}}$ | $x_{\text{f}}$ | $\frac{1}{1 + e^{(v+78)/5}}$ | $\frac{1900}{1 + e^{(v+15)/10}}$ | | $\frac{\alpha_{x_{\text{f}}}}{(Q_{10}^{\text{f}})^{Q_p}}$ |

**Figure S2:**
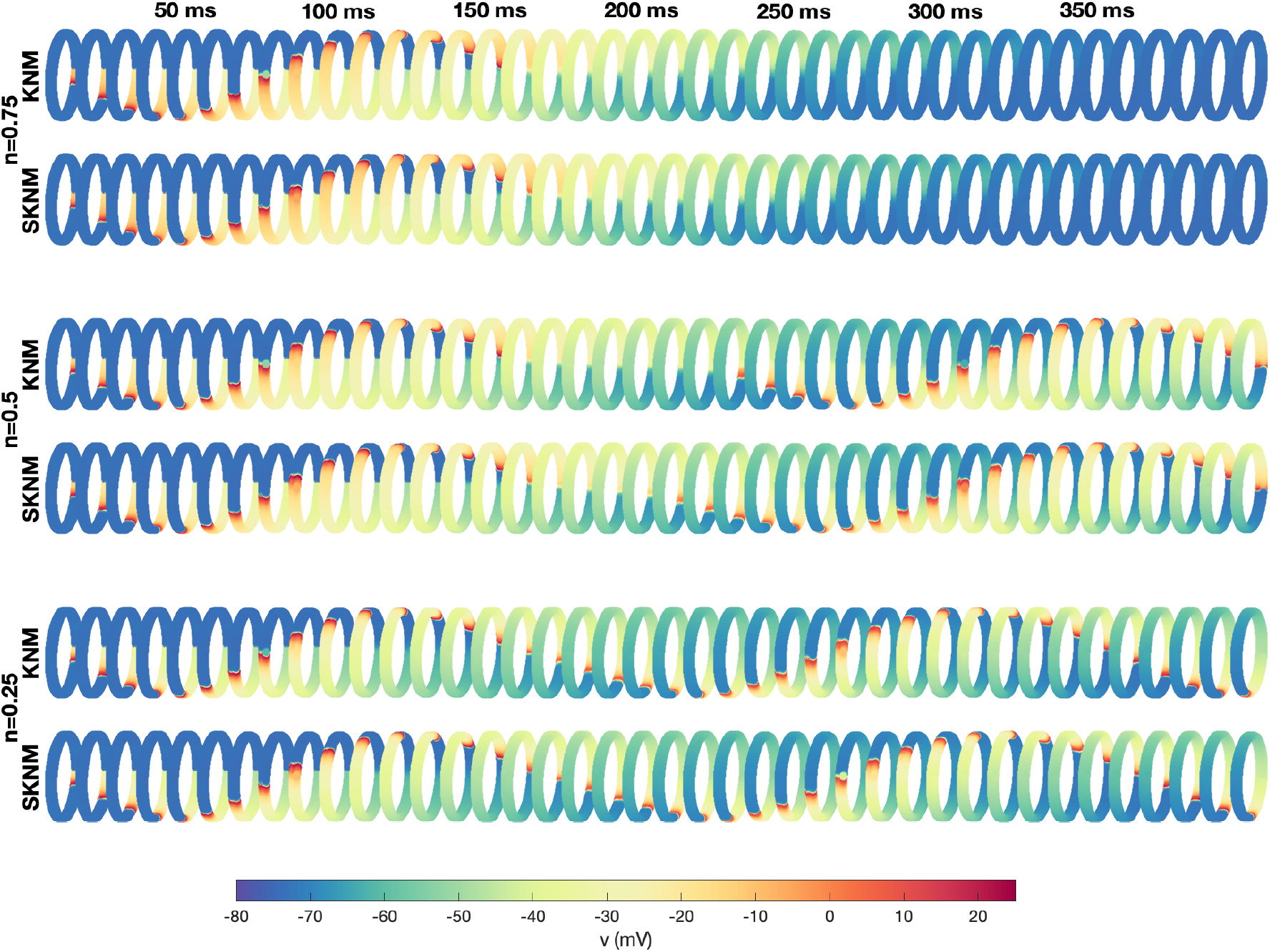
Comparison between KNM and SKNM solutions. The figure shows the membrane potential for the cells in the PV sleeve cylinder at some different points in time after a stimulation is applied for some KNM and SKNM simulations of *n* = 0.75, *n* = 0.5 and *n* = 0.25. We observe that the two models produce quite similar solutions, but that the conduction velocity does not seem to be entirely identical for the two models.

**Figure S3:**
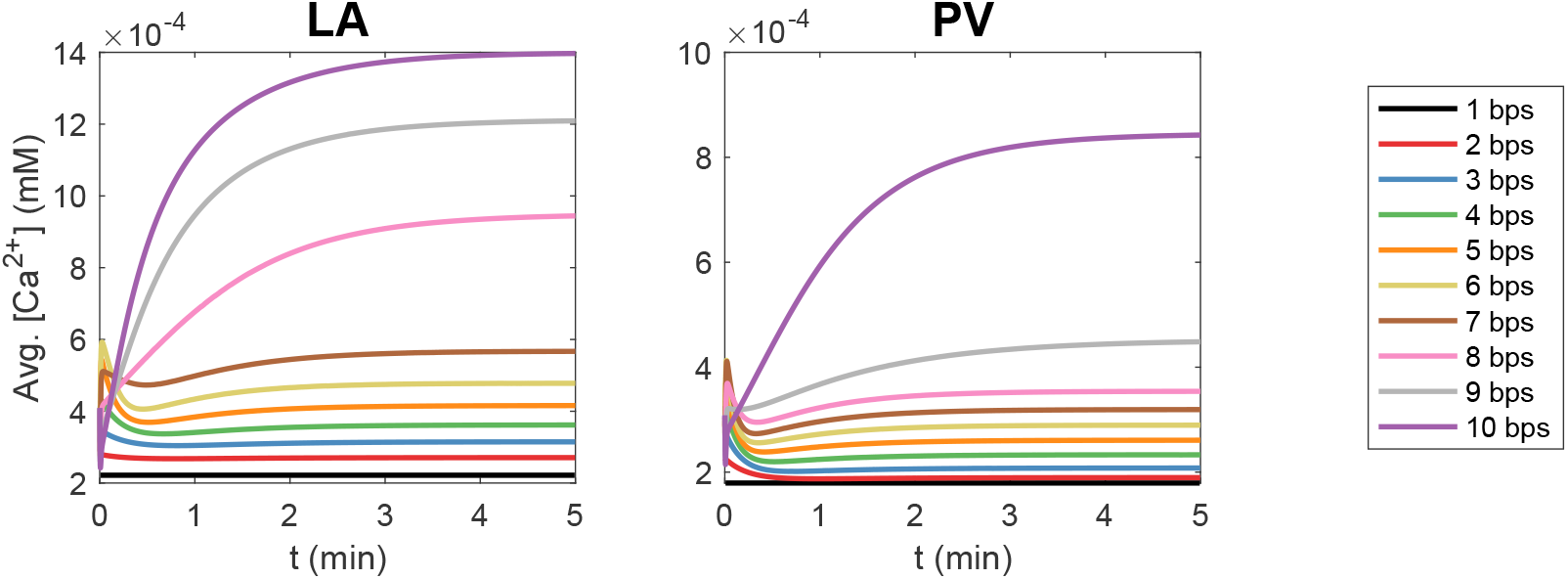
Effect of rapid pacing on the intracellular calcium concentration in the PV and LA membrane models without protein regulation. The figure shows the time evolution of the average cytosolic calcium concentration as a function of time for pacing frequencies ranging from 1 beat per second (bps) to 10 bps.

#### S3.1 Effect of rapid pacing on the cytosolic calcium concentration

In Figure S3, we investigate the effect of the pacing frequency on the average cytosolic calcium concentration between each stimulation. We consider pacing frequencies ranging from the default 1 beats per second (bps) up to 10 bps. We observe that after a few minutes of rapid pacing, the cytosolic calcium concentration increases and that the increase in calcium concentration is larger for a higher pacing frequency.

#### S3.2 Effect of adjusting currents on the cytosolic calcium concentration

In Figure S4, we report the percent change in the average cytosolic calcium concentration resulting from a 20% increase in each type of ion channel, pump and exchanger in the cell membrane for the LA and PV membrane models without protein regulation. We observe that increasing the number of channels responsible for the fast sodium current (*I*_Na_), the late sodium current (*I*_NaL_) or the background sodium current slightly increases the intracellular calcium concentration. Furthermore, increasing the channels responsible for the L-type calcium current (*I*_CaL_) or the background calcium current (*I*_bCa_) results in a considerable increase in the intracellular calcium concentration. On the other hand, increasing the number calcium pumps (*I*_pCa_), sodium-calcium exchangers (*I*_NaCa_), sodium-potassium pumps (*I*_NaK_), background chloride channels (*I*_bCl_), or *I*_Kr_, *I*_Ks_, *I*_to_, or *I*_Kur_ potassium channels, causes an decrease in the cytosolic calcium concentration. The effect of increasing the number of *I*_K1_ and *I*_f_ channels on the cytosolic calcium concentration is small, and differs between the PV and LA versions of the model.

**Figure S4:**
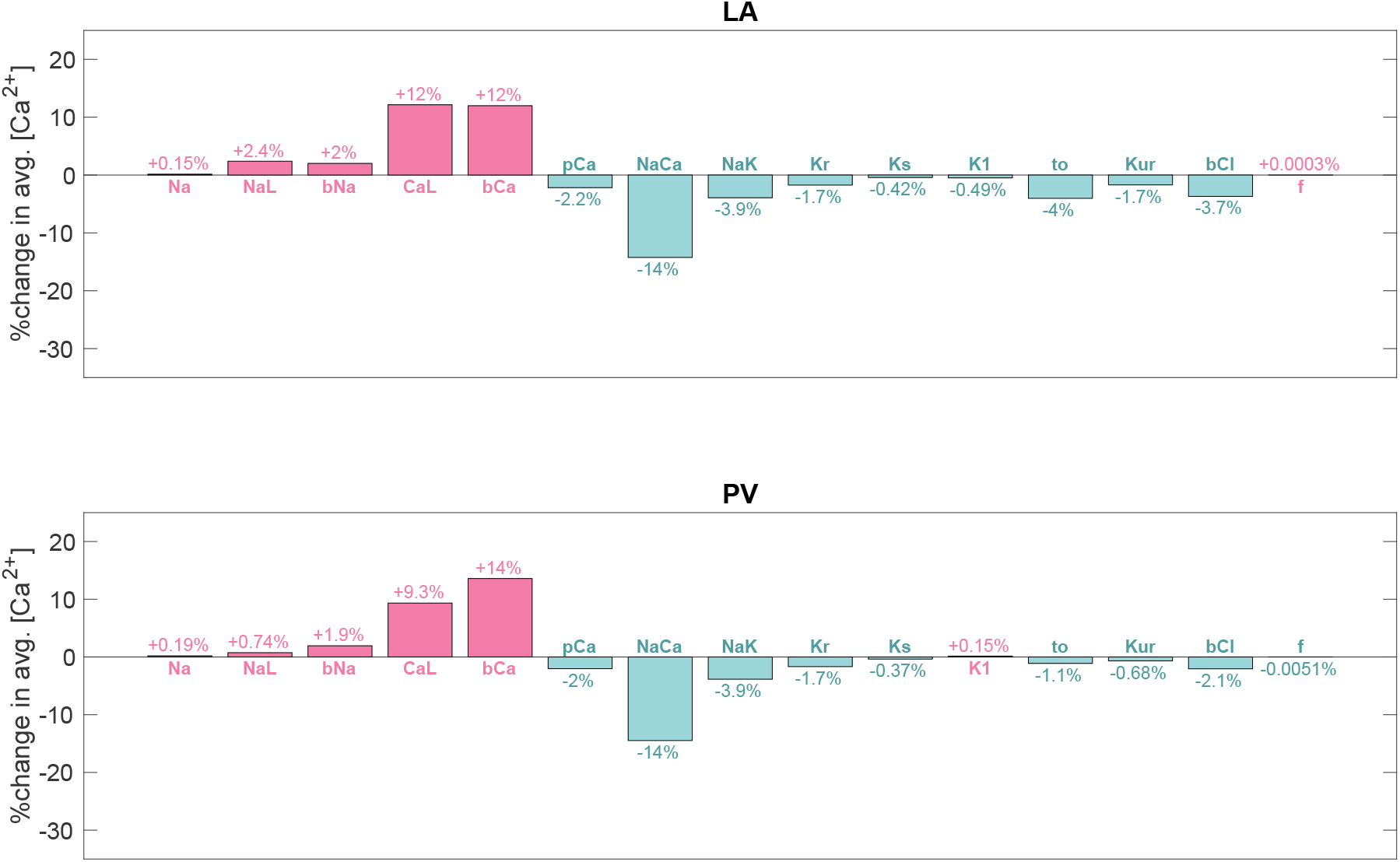
Effect on the intracellular calcium concentration of increasing the number of different types of membrane proteins. For the LA and PV membrane models without protein regulation, we report the percent change in the average cytosolic calcium concentration resulting from a 20% increase in each type of ion channel, pump and exchanger in the cell membrane. The concentrations are recorded 1 h after the parameter change was applied.

